# Progress toward the identification and stacking of crucial domestication traits in pennycress

**DOI:** 10.1101/609990

**Authors:** Ratan Chopra, Evan B. Johnson, Ryan Emenecker, Edgar B. Cahoon, Joe Lyons, Daniel J. Kliebenstein, Erin Daniels, Kevin M. Dorn, Maliheh Esfahanian, Nicole Folstad, Katherine Frels, Michaela McGinn, Matthew Ott, Cynthia Gallaher, Kayla Altendorf, Alexandra Berroyer, Baraem Ismail, James A. Anderson, Donald L. Wyse, Tim Umaslov, John C. Sedbrook, M. David Marks

## Abstract

The oilseed species *Thlaspi arvense* (pennycress) is being domesticated as a new crop that can provide both important ecosystem services and intensify farmland output. Through the use of high throughput sequencing and phenotyping, along with classical mutagenesis key traits needed for pennycress domestication have been identified. Domestication traits identified herein include reduced pod shatter, early maturity, reduced seed glucosinolate levels, and improved oil fatty acid content. By taking advantage of pennycress’ close genetic relationship with *Arabidopsis thaliana*, the causative mutations responsible for each of these traits have been identified. These mutations have been used to develop molecular markers to begin to stack the traits into individual lines.

---

Tremendous gains in crop yields have been achieved through a combination of increased fertilizer use, crop breeding, and improved agronomic practices (2014). However, the long-term sustainability of crop production using current farming practices is threatened by several factors. In intensive cropping systems, such as the corn-soybean rotation in the Midwestern United States, only 40-60% of the applied nitrogen fertilizer is taken up by plants (Wortmann *et al.* 2011). Much of the remaining nitrogen is released as greenhouse gases or pollutes surface and ground waters resulting in the eutrophication of streams and lakes, the creation of dead zones in coastal regions, and the contamination of wells in rural communities (Robertson and Vitousek 2009). Another looming threat to agricultural production is the emergence of herbicide tolerant weeds, including weeds tolerant to glyphosate (Heap 2014). The use of glyphosate resistant crops over the past twenty-five years enabled no-till or reduced-till agriculture that helped to reduce soil erosion (Price *et al.* 2011). Due to the increasing prevalence of herbicide resistant weeds farmers may return to deep tillage to reduce weed pressures, which will hasten soil erosion (Price*, et al.* 2011).

To confound the above issues, the world population is expected to grow to over nine billion by the year 2050 (Gerland *et al.* 2014). To feed this growing population, it has been estimated that food production must increase by an average of 44 million metric tons per year (Tester and Langridge 2010). In addition, global warming due the release of CO_2_ from fossil fuels is posing another threat (Anderson *et al.* 2016). To reduce this threat demand is increasing for plant-based renewable feedstocks for the production of biofuels and bioproducts (Ho *et al.* 2014). However, this leads to the concern that using traditional crops for biofuel and bioproducts formation will threaten future food security (Harvey and Pilgrim 2011). One solution is to intensify farmland output (Tilman *et al.* 2011).

The deployment of cover crops during the fallow period between the growth of summer crops can improve water quality and reduce the threat of herbicide tolerant weeds (Dunn *et al.* 2016). However, traditional cover crops such as winter rye often need to be terminated before maturity to allow the planting of summer crops (Roesch-McNally *et al.* 2017). Thus, such cover crops do not enhance food or feedstock production. The winter annual weed, pennycress (*Thlaspi arvense* L. also known as field pennycress and referred to as “pennycress” hereafter), is being domesticated as a new alternative cover crop (DeHaan *et al.* 2016, Isbell 2009, Jordan *et al.* 2007, Phippen and Phippen 2012, Sedbrook *et al.* 2014). As a winter cover, pennycress can utilize excess nitrates before they escape into the environment and can suppress the growth of spring weeds (Johnson *et al.* 2015, Johnson *et al.* 2017, Weyers *et al.* 201). Importantly, pennycress can be harvested for its oilseeds (Fan *et al.* 2013, Moser 2012, Moser *et al.* 2009a, Moser *et al.* 2009b). Thus, pennycress has the potential to intensify farm output by producing a new crop on land that is temporally held fallow, such as much of the tens of million hectares of land currently undergoing the corn/soybean rotation in the Midwestern United States (Hart 2015).

The selection of pennycress as a target for domestication as a new cover crop was based on positive natural traits such as extreme winter hardiness (−25°C), high seed yields for a wild species (1,000-2,000 kg/ha), and seeds rich in oil (30-35%) and protein (25-27%) (Warwick *et al.* 2002). Pennycress produces an oilseed suitable for making biofuels and bioproducts (Fan*, et al.* 2013, Moser 2012, Moser*, et al.* 2009a). However, the oil and associated seed meal are not suitable for human or animal consumption. The oil is enriched with erucic acid, which has been considered to be unsuitable for human consumption(Bell 1982). In addition, the seeds contain glucosinolates, which are considered to be anti-nutritional for humans and non-ruminant animals (Wittkop *et al.* 2009). Furthermore, the mature seedpods are prone to breakage or shatter resulting in pre-harvest seed loss. Lastly, in many regions, pennycress matures late enough to cause delays in the planting of summer crops. Successful domestication of pennycress requires the elimination of these negative traits.

Herein, we show that the rapid domestication of pennycress can be facilitated by its close relationship to the model plant Arabidopsis (Sedbrook*, et al.* 2014). Like Arabidopsis, pennycress is self-fertile and has a relatively small non-repetitive diploid genome. Genome sequencing and analyses revealed a mostly one-to-one functional correspondence between single *Arabidopsis* genes and single pennycress ortholog genes (Chopra *et al.* 2018b, Dorn *et al.* 2013, Dorn *et al.* 2015). Importantly and directly related to the domestication of pennycress, recessive Arabidopsis mutants have been described that eliminate weedy and other agronomically undesirable traits (Provart *et al.* 2015, Sedbrook*, et al.* 2014). The goal of this study was to identify similar mutants in pennycress with traits considered to be crucial to the domestication of any plant species (Abbo *et al.* 2014). Using classical mutagenesis, we describe the isolation and characterization of pennycress mutations conferring crucial domestication traits such as early flowering/maturity, reduced seedpod shatter, reduced glucosinolates, and reduced erucic acid along with reduced polyunsaturated fatty acids resulting in the creation of canola-like high oleic oil. The causative mutations responsible for each of these traits were identified. These mutant sequences were used to develop molecular markers that have been successfully employed to stack all of the mutations into a single line.

## Results

### Early flowering/maturing pennycress

For pennycress to fit between the rotations of traditional summer crops it needs to mature in the spring without disrupting the planting of the following summer crop (De Bruin and Pedersen 2008). To identify Arabidopsis-like mutants that flower and mature early, pools of mutagenized population of M_2_ seeds were planted in the field space adjacent to the University of Minnesota St Paul campus (Chopra*, et al.* 2018b). During the following spring, plants in the field that flowered early were tagged and followed through to maturity. Sixty-one independent isolates that flowered and matured early compared to the wild-type were identified during the primary screen. Nineteen of the 61 early flowering mutants were confirmed for the inheritance of earliness traits in the following generation. We further focused our efforts on one line, A7-25, that flowered and began producing seedpods 10-14 days earlier than the wild-type (compare Figs. 1a, b). In another site-year in St. Paul, this mutant line matured approximately one week ahead of the wild-type allowing an early harvest in the field conditions while demonstrating normal plant stature and yield (Table. S1)

**Figure. 1.**
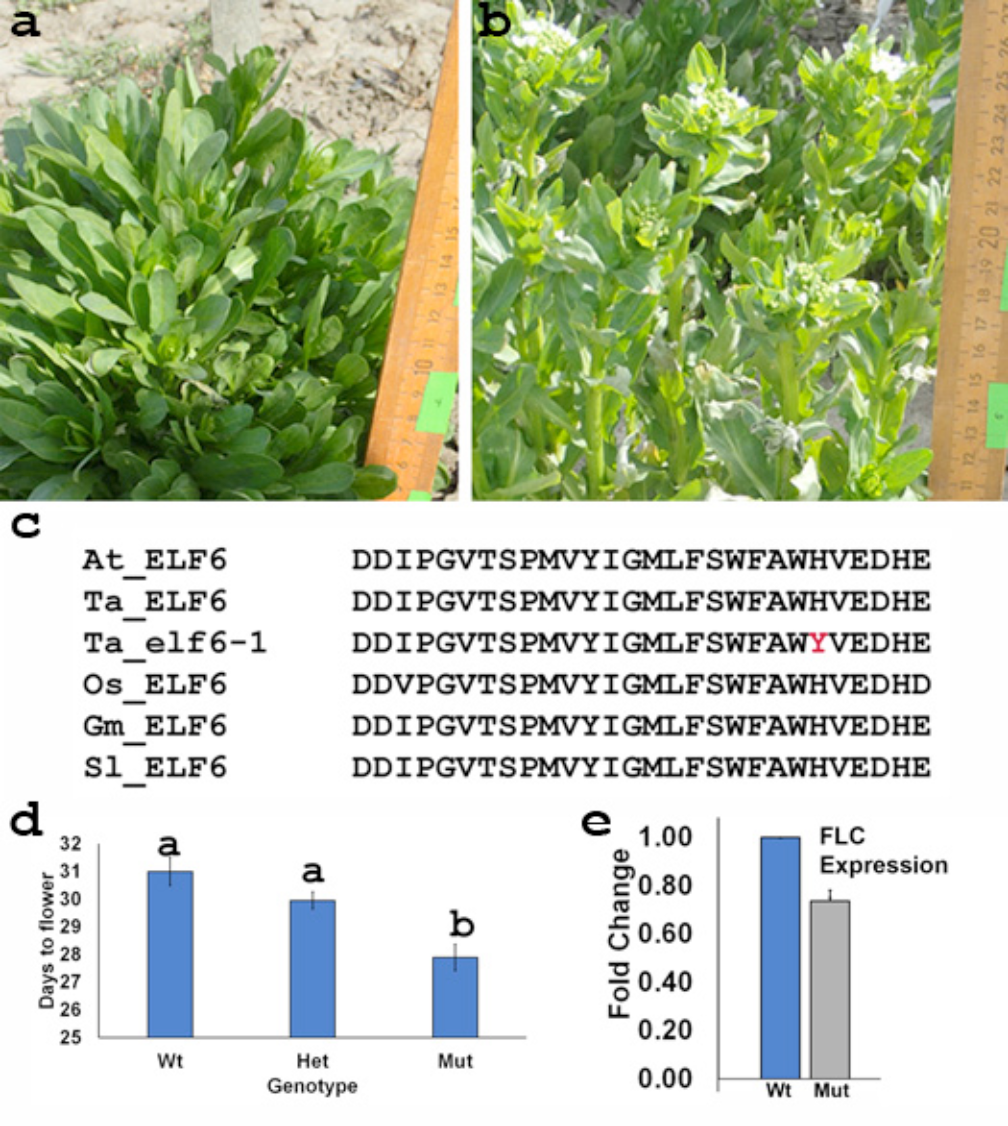
Early flowering pennycress mutant. Images of field grown wild-type (**a**) and early flowering *Ta-elf6-1* mutant (**b**) plants taken on the same date. (c) Comparison of amino acid sequences within the JmjC domain of *ELF6*-like genes highlighting the residue altered in the pennycress mutant (red). (**d**) Average days to flowering in greenhouse conditions for an F_2_ population derived from a cross between wild-type and *Ta-elf6-1* plants. Allele-specific markers were used to assess the *elf6-1* genotype of members in the F_2_ population. Note: Letters indicate significant differences based on pair-wise Tukey test. (**e**) qPCR analysis of *FLC* expression in wild-type and early flowering *Ta-elf6-1* mutant plants using RNA from field grown plants collected in the fall of 2017. *FLC* expression values were normalized using a ubiquitin probe (error bar denotes standard deviation). Abbreviations: At – *Arabidopsis thaliana*, Os – *Oryza sativa* (rice), Gm – *Glycine max* (soybean), Sl-*Solanum lycopersicum* (tomato).

To test the hypothesis that the A7-25 mutant carried a mutation in a previously characterized in Arabidopsis gene, whole-genome sequencing (WGS) was performed. It was found that A7-25 contains a mutation in the candidate pennycress ortholog of the Arabidopsis *EARLY FLOWERING 6* (*ELF6*) (referred to as *Ta-ELF6-1* in Fig. 1c, and Data S1-3) (Jeong *et al.* 2009, Noh *et al.* 2004). In Arabidopsis, ELF6 activity de-represses the expression of the floral inhibitor *FLOWERING LOCUS C* (*FLC*) gene by removing methyl groups from lysine 27 of histone 3 proteins associated with the *FLC* locus (CrevillÃ©n *et al.* 2014). *FLC* controls flowering time in pennycress(Dorn *et al.* 2017). The reduced demethylase activity in *elf6* mutants results in reduced *FLC* expression, which hastens flowering. As shown in Fig. 1c and Data S3 the mutation in *Ta-ELF6-1* results in the substitution of a tyrosine (Y) residue for a conserved histidine (H) (Noh*, et al.* 2004). This substitution is predicted to disrupt the formation of the iron-binding site in the Jumonji C (JmjC) domain that is required for histone 3 demethylase activity in many plant species(Lu *et al.* 2008). To determine if the early flowering phenotype in pennycress co-segregates with the mutation, an F_2_ population derived from a cross between the mutant and wild-type (Ames23761) was characterized. Statistical analysis of each genotype class in the F_2_ population showed that plants homozygous for the *Ta-elf6-1* mutation flowered earlier than either homozygous wild-type or heterozygous plants (*p*-values 0.0002 and 0.0021, respectively (Fig. 1d).

All studies with Arabidopsis on the effects of the *elf6* mutation on *FLC* expression have been under controlled growth conditions. To determine if a similar phenomenon occurs in field grown pennycress, we analyzed the effect of the mutation on pennycress *FLC* expression prior to vernalization in the fall. As shown in (Fig. 1e), *FLC* expression was reduced by 30.5±0.09% in the early flowering A7-25 line compared to the wild-type grown in the same trial. These results show that *ELF6* functions the same under field conditions for pennycress as under controlled growth conditions for Arabidopsis.

### Reduced seedpod shatter pennycress

Pennycress seedpods exhibit premature breakage by wind or during mechanical harvest resulting in yield losses of over 50%. To identify pennycress mutants with reduced seedpod shatter, fields of M_2_ plants maintained past peak maturity were screened for individuals exhibiting reductions in pod breakage or shatter (compare Figs. 2a, b). Five individual recessive mutants have been identified, along with a sixth line identified in a growth room screen, in which the trait successfully passed through multiple generations.

**Figure 2.**
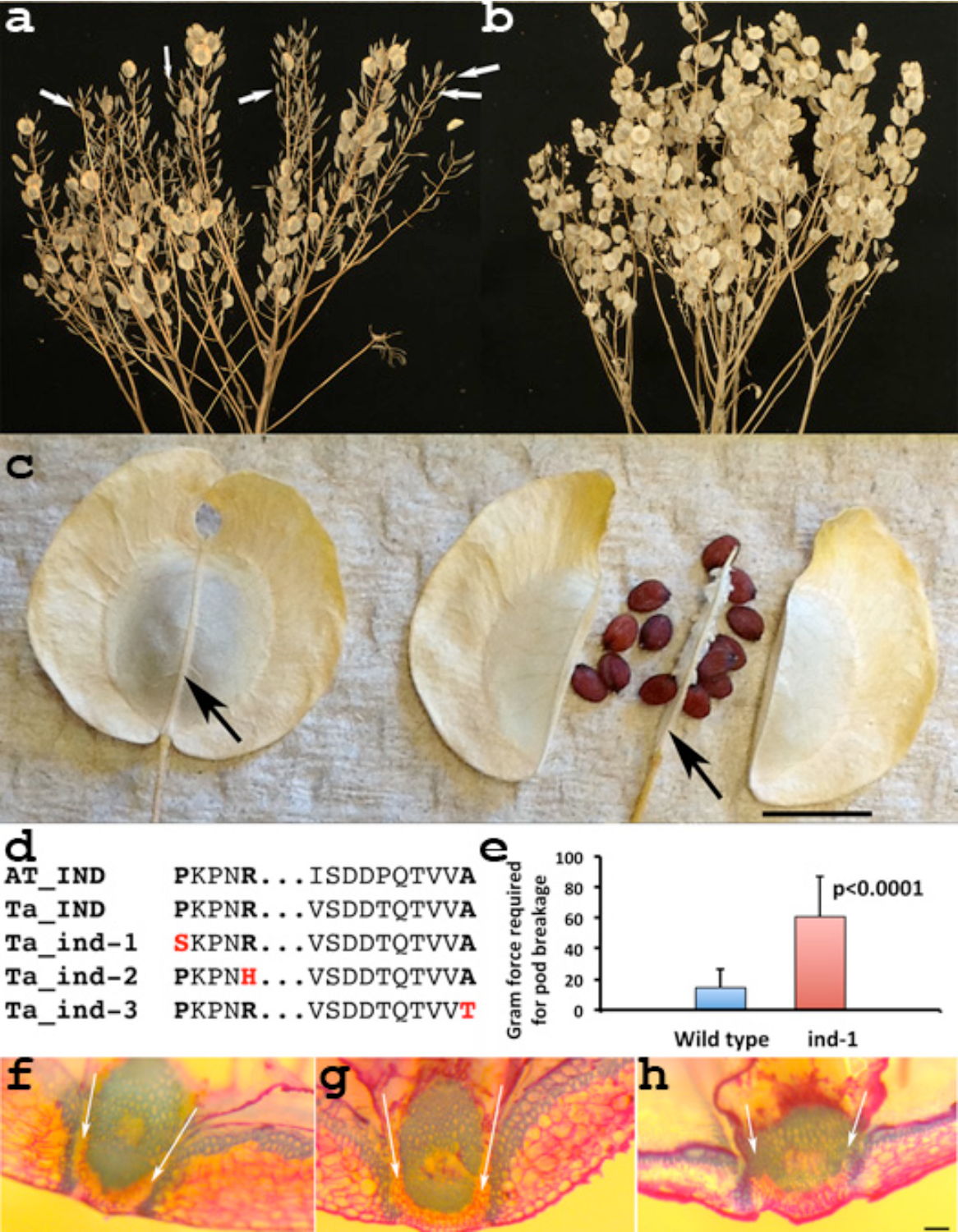
Isolation and characterization of pennycress seed pod mutants. (**a**) Field grown wild-type plant compared to a reduced shatter mutant plant (**b**). (**c**) An intact seedpod compared to a broken seedpod with released seeds (both are wild-type). The arrows show the septum of an intact pod (left) and the septum structure that remains after pod shatter. The arrows highlight the septum remaining after pod shatter. (**d**) IND amino acid sequences derived from Arabidopsis (At-IND) and wild-type pennycress (Ta-IND) compared to three mutant pennycress alleles (*ind-1*, *ind-2* and *ind-3*). The red letters highlight the amino acid substitutions. (**e**) Gram force needed to break open pods from wild-type and *ind-1* plants. Std. dev. error bars are shown. Freehand sections through wild-type (**e**), *ind-1* (**f**), and *ind-2* (**g**) seedpods. The arrows highlight the separation layers in wild-type and *ind-1*. This layer is missing in *ind-2* and *ind-3* (Fig. S2). The darker stained blue regions highlight lignified cell layers. The size bars in **c** and **h** represent 0.5 cm and 50 µm, respectively.

Pennycress produces a flat circular seedpod called a silicle that has a visibly distinct morphology compared to the long rod-shaped seedpods known as siliques produced by Arabidopsis and other members of the Brassicaceae plant family (Fig. 2c). Therefore, it was unknown if genes required for pod formation in Arabidopsis would be the same as those controlling pod formation in pennycress. To determine if the pennycress mutants contained mutations in genes known to be associated with the formation of the separation zone in Arabidopsis, which mediates pod breakage, the six lines were subjected to WGS. Three of the lines were found to have mutations in the pennycress candidate ortholog of the Arabidopsis *INDEHISCENT* (*IND*) gene (Fig. 2d, Data S1–3) (Liljegren *et al.* 2004). The analysis of the remaining three lines will be considered elsewhere. *IND* encodes a *bHLH* transcription factor that regulates the formation of separation layers of cells contained within the value margin, which flanks the septum (also called a replum) that divides the seedpod into to two halves (Liljegren*, et al.* 2004). The *Ta-ind-1* allele contains a mutation outside of a highly conserved bHLH domain, while the *Ta-ind-2* flanks this region and *Ta-ind-3* has a mutation in the conserved bHLH domain (Fig. 2d and Table S2). The seedpods of both *Ta-ind-2* and *Ta-ind-3* failed to cleanly break on harvest signifying non-functional dehiscence zones. However, *Ta-ind-1* seedpods did break at the septum to release seeds on harvest, but compared to wild-type, required more force to break open the seedpods (Fig. 2d). As expected, *Ta-ind-1* retains a separation layer (Fig. 2e) that is not present in *Ta-ind-2* or *Ta-ind-3* (Fig. 2f and Fig. S1). The finding of three independent pennycress *ind* alleles showing reduced pod breakage strongly supports the causation of these mutations of the observed phenotype. Of the three alleles, *Ta-ind-1* should have the most agronomic utility, as the shatter-less phenotype of *Ta-ind-2* and *Ta-ind-3* reduces seed harvestability with conventional combines.

### Low glucosinolate pennycress

Like Arabidopsis and other brassicas, pennycress seeds contain glucosinolates (~100 µmol/g of seeds), almost all in the form of sinigrin (Warwick*, et al.* 2002). Most of the glucosinolates remain in the meal after the extraction of oil from the seeds. This seed meal has added value if it can be used as an animal feed supplement, however high amount of glucosinolates in the meal can negatively affect animals (Wittkop*, et al.* 2009). Thus, a search for pennycress lines with reduced glucosinolates was conducted. As described in a previous report, 15,000 lines were scanned using Near Infra-Red Spectroscopy (NIRS) to estimate seed composition traits in the intact seeds (Chopra *et al.* 2018a). The analysis of these scans identified one line predicted to have very low glucosinolate levels, which was confirmed via wet lab assays for total seed glucosinolates (Fig. 3a). Interestingly, numerous individuals indicated that the mutant seeds have a pleasant, nutty flavor that has not previously been reported for seeds from plants in the Brassicaceae plant family. Seeds from the Brassicaceae family typically are characterized to have flavors that range from bitter, mustardy, garlicky, to hot. Hereafter, this pennycress mutant is referred to as “nutty”. The leaves of nutty also lacked the garlicky smell that accounts for pennycress also being called “stinkweed”.

**Figure 3.**
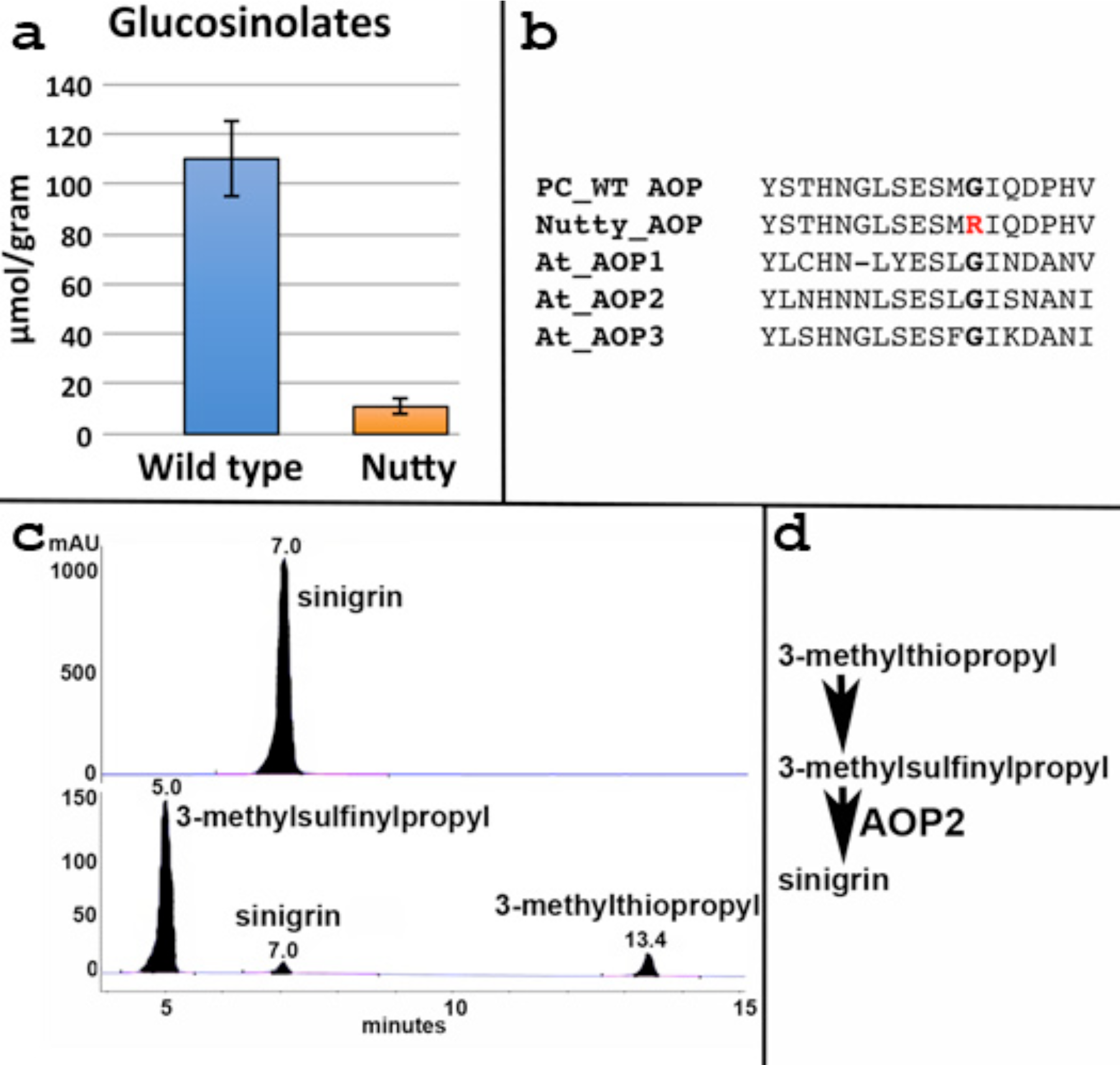
Comparison of glucosinolates in wild-type and Nutty (*aop2-1*) mutant pennycress. (**a**) Spectrometric quantitation of total glucosinolates in wild-type and the nutty mutant. Error bars highlight Std. Dev. (**b**) AOP-like amino acid sequences derived from Arabidopsis (At-AOP 1 through 3) and wild-type pennycress AOP2 (PC_WT AOP) compared to the nutty mutant (Nutty-AOP). (**c**) HPLC analysis of desulfated glucosinolates from wild-type and the nutty mutant, top and bottom, respectively. Peaks are labeled with compound name and the time (minutes) of flow through the HPLC. (**d**) Terminal biochemical steps leading to the synthesis of the glucosinolate sinigrin. The step mediated by the enzyme encoded by *AOP2* in Arabidopsis is shown.

In Arabidopsis, the biochemical pathway leading to the formation of various glucosinolates including sinigrin has been well characterized(Sonderby *et al.* 2010). Thus, we conducted WGS on nutty and searched for a mutation in a gene known to be involved in glucosinolate biosynthesis in Arabidopsis. This led to the identification of a pennycress mutation in a gene that is equally related to a family of three tandemly linked genes in Arabidopsis called *ALKENYL HYDROXALKYL PRODUCING 1, 2, and 3* (*AOP1,2,3*)(Kliebenstein *et al.* 2001). The analysis of an F_2_ from a cross between wild type and mutants showed that the reduced glucosinolate nutty phenotype was tightly linked to the mutation in the AOP-like sequence (Fig. S2). HPLC analysis of glucosinolates from nutty confirmed the reduction in sinigrin and also revealed a mild excess accumulation of 3-methylsulfinylpropyl and 3-methylthiopropyl glucosinolates, which are precursors to sinigrin in the biosynthetic pathway, supporting the hypothesis that the pennycress *AOP-like* gene encodes an enzyme with *Arabidopsis AOP2* alkenyl producing activity converting 3-methylsulfinylpropyl glucosinolate to sinigrin (Fig. 3d) *(Kliebenstein, et al. 2001).* Thus, the mutant gene was named *Ta-aop2-*1. Of note, all three glucosinolates shown in the HPLC profile shown in Fig. 3c can be detected in the glucosinolate profile shown in Fig. 3a. If there was a simple blockage in the pathway one would predict that the total levels of glucosinolates would be the same in both mutant and wild type, however, there is a clear reduction in glucosinolates in nutty. Additional work is needed to further understand the molecular mechanisms responsible for the overall reduction in glucosinolates in this mutant.

### Improving fatty acid composition of pennycress

The fatty acid composition of pennycress oil is similar to that of rapeseed containing over 35% erucic acid (Fig. 4a) (Claver *et al.* 2017, Moser*, et al.* 2009a). Oils containing such high levels of erucic acid are not considered fit for human or animal consumption(Knutsen *et al.* 2016). Therefore, a search was made for lines with reduced erucic acid. This search was aided by the knowledge that in Arabidopsis and other species, the biosynthesis of erucic acid is mediated by a single enzyme, *FATTY ACID ELONGATION1* (*FAE1*), which in two steps converts oleic acid with eighteen carbons and one double bond (18:1) to erucic acid with twenty-two carbons and one double bond (22:1) (James *et al.* 1995). Similar to the detection of glucosinolates, the quantity of erucic acid along with other types of fatty acids can be estimated using NIRS(James*, et al.* 1995). Thus, the same 15,000 NIRS scans that were used to identify the reduced glucosinolate line were used to search for lines with reduced erucic acid. This analysis identified two independent lines with greatly reduced erucic acid. These results were verified via gas chromatography (GC), which showed that both lines only contained trace levels of erucic acid (Fig. 4b). The candidate pennycress *FAE1* alleles in these reduced erucic acid lines were amplified via PCR and sequenced. Both lines contained independent nonsense mutations leading to premature stops in the predicted pennycress FAE1 protein sequences (Data S1–3). These results were further confirmed by WGS of these lines. The finding of two strong loss of function alleles for the candidate pennycress *FAE1* ortholog giving rise to the expected phenotypes strongly supports causation of the mutations.

**Figure 4.**
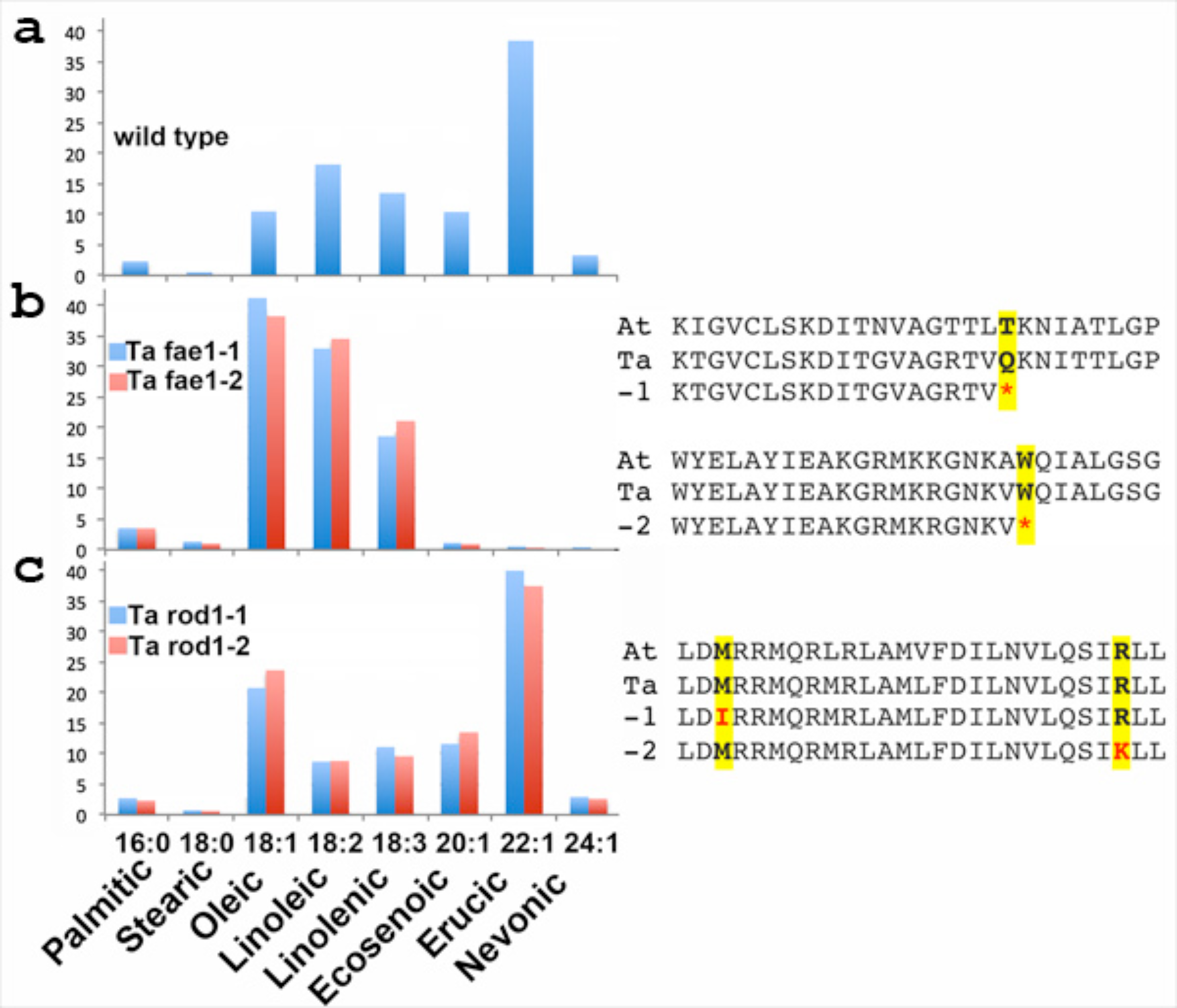
Comparison of wild type fatty acid profiles compared to those of pennycress *fae1* and *rod1* mutant lines. (**a-c**) Profiles of wild-type pennycress, pennycress *fae1* and *rod1* mutants. The amino acid sequences derived from Arabidopsis and wild-type pennycress compared to mutant pennycress alleles are shown to the right. The red letters highlight the amino acid substitutions. Fatty acid names along with carbon chain lengths and the degrees of desaturation are shown across the bottom.

Pennycress *fae1* mutants contain the desired reduction in erucic acid and a desirable elevation in oleic acid. They also accumulate higher levels of fatty acid with extra double bonds referred to as polyunsaturated fatty acids (PUFAs). In particular, linolenic acid with 3 double bonds (18:3) was elevated. This fatty acid belongs to the omega3 class of fatty acids, which are associated with reduced heart disease, reduced bone fracture risk, and in reduced childhood obesity (Perng *et al.* 2014, Rajaram 2014). However for many applications, PUFAs are not desirable as the extra double bonds reduce the stability of the oil, thus shortening the shelf life of the products that contain the oil (Gordon 2001). Therefore, a search was made for lines containing reduced levels of PUFAs using the same NIRS screen. We identified two mutants with predicted reductions in both linoleic and linolenic acids. These reductions were verified using GC analysis (Fig. 4c). Both lines contained similar levels of erucic acid as the wild-type, but the reductions in PUFA were compensated for by an increase in oleic acid. The fatty acid profiles of these two lines resembled those from Arabidopsis *reduced oleate desaturation1* (*rod1*) mutants(Lu *et al.* 2009). This was confirmed via the PCR amplification and sequencing of the candidate pennycress genes (**Data S1-3**). These results were further confirmed by WGS of these lines.

### Stacking traits to develop domesticated pennycress

This study has identified pennycress lines that harbor crucial domestication traits including early maturity, reduced shatter, reduced erucic acid and reduced PUFAs. Importantly, none of the described mutations reduced plant stature or predicted yield (Table S1 and Fig. S3). Knowledge of the causative mutations in each of the lines has been used to develop molecular probes that can follow each of the traits in segregating populations at the molecular level (Table S3-4 and Data S1–3). To begin to combine these traits, a cross was made between *Ta*-*fae1-1* and *Ta*-*rod1-1*. KASP markers were designed for the specific mutations in each of these mutants (see Materials and Methods). Using this approach (Fig S4) it was possible to identify the double mutants in an F_2_ population at the seedling stage over two months before they began to produce seeds. The fatty acid composition of the oil from these double mutants was very high in oleic acid, low in erucic acid, with reduced levels of in PUFAs. Overall, the fatty acid profile of the double mutant pennycress oil closely resembled the profile of canola oil (**compare** Figs. 5a and 5b) (Dyer *et al.* 2008).

**Figure 5.**
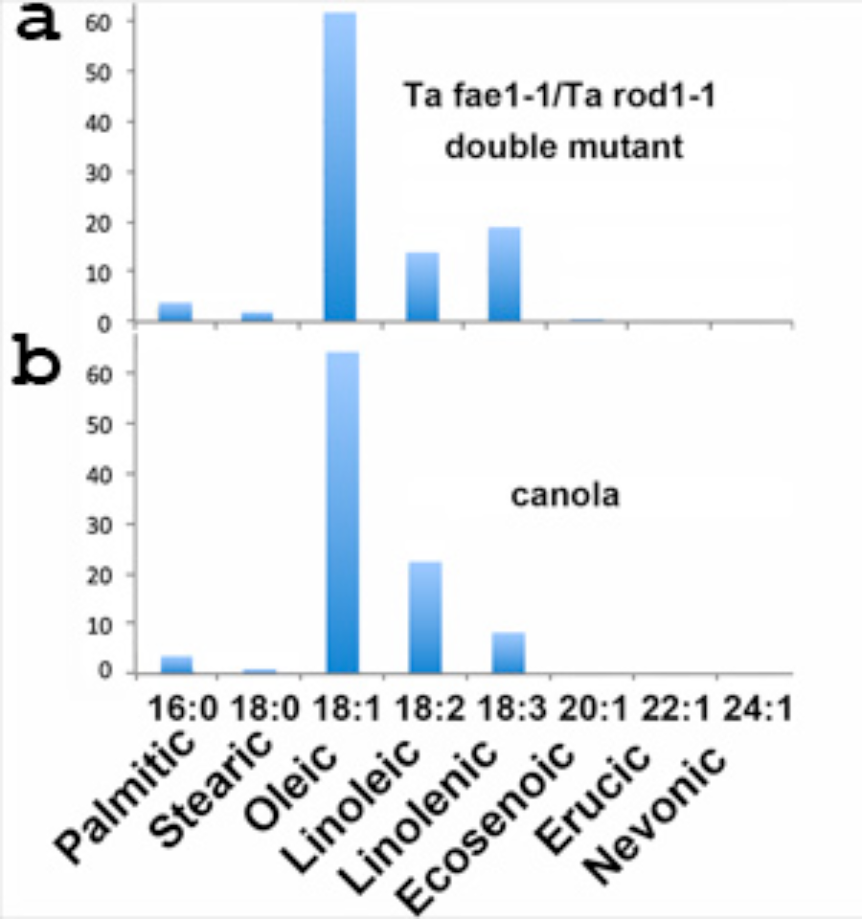
Comparison of fatty acid profiles of pennycress fae1/rod1 double mutant compared to canola oil. (**a**) Double mutant, (**b**) canola. Fatty acid names along with carbon chain lengths and the degrees of desaturation are shown across the bottom.

Mutation specific KASP markers have been used to identify additional double and triple mutant combinations in F_1_ and F_2_ individuals derived from various crosses between the pennycress mutants (Table S5). This includes a line that is homozygous for the combined *Ta-fae-1*, *Ta-rod-1*, and *Ta-aop2-1* mutations. This line produces seeds that are essentially equivalent to those from the so-called “double 0” canola that similarly lack erucic acid and contain low levels of glucosinolates(Bell 1982). Additional crosses have generated an F_1_ plant that is heterozygous for all five of the desirable traits described in this report (Fig S5). Progenies from this F_1_ plant will be genotyped to identify a line homozygous for all five mutations. This stacked line will represent the first-generation domesticated pennycress variety and will be subjected to field-testing ahead of potential commercialization.

## Discussion

The traits described in this report should allow domesticated pennycress to intensify farm output by utilizing land from fall to spring that is not currently in production. The early maturing pennycress described in this report will better fit into existing crop rotations. The reductions in pod shatter will greatly improve harvestable yield, and the reductions in glucosinolates and improvements in oil quality will enhance the value of the oilseeds. The rapid progress in identifying these traits in pennycress was due to reduced gene redundancy, which facilitated the creation of pennycress mutants that closely resemble similar Arabidopsis mutants(Chopra*, et al.* 2018b, Sedbrook*, et al.* 2014).

The causative mutations responsible for the domestication traits described in this study have been well studied in *Arabidopsis*(Provart*, et al.* 2015). Knowing the causative nature of the mutations responsible for the pennycress domestication traits is important for several reasons. The information can be used to stimulate additional basic research on the molecular mechanisms that control these traits. This is especially true for the *Ta-AOP2* gene. The enzyme encoded by this gene catalyzes the last step in the glucosinolate pathway in pennycress. Through some unknown mechanism the mutation in this gene results in reduced metabolic flux through the pathway. The information also can be used to improve current crops or to help guide the domestication and creation of new crops. For example, *ELF6* now becomes good target for hastening maturation of related *B. napus* (canola). Likewise these results show that simultaneously reducing in both *ROD1* and *FAE1* activity can greatly increase oleic acid content above that found for either of the single mutants. This should have wide ranging applications in modifying the seed oil fatty acid content of many plants. Importantly, the causative mutations can be used as molecular markers as in the pipeline shown in Fig. S4 to combine the pennycress traits into single lines. Using this pipeline, single plant lines harboring all the traits described in this report will be ready for field-testing within the next two years.

The traits described in this report were created using classical mutagenesis. Classical mutagenesis has long been used to induce useful mutations in crop plants (Ahloowalia *et al.* 2004, Oladosu *et al.* 2016). For this work we used EMS as a mutagen (Chopra*, et al.* 2018b). Treatments with EMS generate up to thousands of mutations in each mutagenized line creating a large pool of mutant genes in a relatively small population. This increases the odds of finding mutants of interest as described in this study. As previously described, these pools mutations can be used to mine for additional desirable traits (Chopra*, et al.* 2018b). On the downside these lines may carry deleterious mutations. However, many of these can be identified from their known affects in Arabidopsis and can be removed by marker-assisted breeding(Chopra*, et al.* 2018b).

In theory, it would have been faster to use gene-editing approaches to create the domestication traits described in this report. In fact, we have created an alternative zero erucic acid pennycress line using CRISPR-Cas9 mediated gene editing (McGinn *et al.* 2018). However, successful utilization of gene editing requires precise knowledge about the genes one wishes to target. This knowledge is not always available. For example, *ELF6, ROD1* and *AOP2-like* were not on the original list of gene targets for improving pennycress(Sedbrook*, et al.* 2014). Furthermore, the creation of partial loss of function mutations, such as that responsible for the desirable *Ta-ind-1* allele, requires prior knowledge before gene editing can be employed. To balance speed and discovery, we will continue to utilize both classical breeding to stack desirable traits and gene editing to produce desirable mutations in existing elite pennycress breeding lines.

There are many obvious improvements that can be made to enhance the potential success of pennycress as a new crop. For example, the dark colored seed coats contain high levels of oxidized tannins, which can interfere with protein uptake in the intestinal tracts of animals that are fed pennycress seed meal(Butler 1992). In a previous report, pennycress mutants lacking tannins have been described, and this trait is currently being added to the stacked line(Chopra*, et al.* 2018b). Additional screens are underway to identify second-generation domestication traits such as reduced seed dormancy, increased yield, increased seed size, increased oil content and improved protein quality. Many of these traits will aid in crop establishment and improve upon current net-economic returns, which increase the likelihood that pennycress will be adopted (Ott *et al.* 2019).

The identification and stacking of domestication traits represent the first step toward the creation a new crop. Concurrently with the ongoing work described in this report, agronomists and ecologists are working to develop planting recommendations and to understand the effects of growing pennycress on the landscape. In addition, food scientists are beginning to study the potential of pennycress as a new food source for human consumption. For example, the “double zero” - like pennycress seeds lacking erucic acid with greatly reduced glucosinolate will be used for various analyses including feeding trials. The canola-like oil will also provide an important new feedstock for the development biofuels and bioproducts.

Newly domesticated pennycress has the potential to intensify farm output by being grown during the fallow period between existing summer crops. Importantly, pennycress grown during the fallow seasons has been shown to provide much needed ecosystem services including nitrogen uptake and the suppression of spring weeds, which will enhance the sustainability of current agricultural practices. We believe the combined oilseed production and ecosystem rewards will greatly outweigh the remaining challenges in developing pennycress as a new cash cover crop.

## Materials and Methods

### Screening of mutant pennycress populations for important traits

We previously described the creation of a large pennycress mutant populations using ethyl methane sulfonate (EMS) and other mutagens (Chopra*, et al.* 2018b). These represent the test populations used to search for pennycress mutants similar to Arabidopsis mutants that exhibit agronomically desirable phenotypes.

### Seed Source, DNA isolation, Sanger Sequencing, Whole-Genome Sequencing

The source of the pennycress seeds for development of genomic resources was previously described in Dorn et al. (Dorn*, et al.* 2015). Mutants characterized in this report were derived from EMS treated seeds as described in Chopra et al.(Chopra*, et al.* 2018b). DNA was isolated from the candidate mutants using a Qiagen plant genomic DNA kit (Qiagen, Valencia, CA). DNA was subjected to WGS using an Illumina HiSeq 2500 sequencer (2×125 bp). Raw reads were analyzed using the method described by Chopra et al.(Chopra*, et al.* 2018b) to detect and predict the nature of mutations in each of these candidate mutants (Table S3). Gene-specific primers (Table S6) were designed to PCR amplify and sequence putative pennycress orthologs. PCR products were subjected to Sanger sequencing. DNA sequences were analyzed using the CLC Genomics Workbench (Qiagen, Valencia, CA) to process files, identify and confirm the corresponding mutations. Protein sequence alignments were performed using the sequences from *Thlaspi arvense* (Ta), and *Arabidopsis thaliana* (At) for the candidate genes. Mutation sites of the candidate genes were changed manually, and alignments were performed using the clustalW program. Genomic DNA sequences, protein sequences and alignments for all the wild-type and mutants genes discussed in this article are provided in the Data S1, S2, and S3 files.

### Early flowering phenotyping

F_2_ plants from a cross between the wild-type GRIN accession Ames 23761 and the early flowering mutant MN A7-25 were planted into Ray Leach SC10 Cone-tainers™ filled with Sun Gro Metro Mix 560 Sun-Coir. After germination, plants were allowed to grow to the two true leaf stage in a 20 ºC growth chamber. Plants were then transferred to a growth chamber maintained at 4 ºC and 8 hours of light for a vernalization period of 21 days. After this period plants were returned to the 20 ºC growth chamber. Days to flowering were recorded as the number of days after plants were returned to the 20 ºC growth chamber to the first open florets visible on the plant.

### Genotyping using allele-specific markers

To perform co-segregation analysis on F_2_ progenies derived from *elf6-1*\*Ames23761 and *aop2-like-1*\*MN106 crosses, we designed allele-specific and flanking primers for each of the alleles (Table S4). DNA was extracted using the Sigma ready extract method and genotyping was performed using the methods described in Chopra et al. (Chopra*, et al.* 2018b).

### Expression analysis

Leaf tissue from ten seedlings for each replicate were pooled from wild-type and the mutant (*elf6-1*) for RNA extractions. RNA was extracted using the RNAeasy mini clean up kit (Qiagen, Valencia, CA) and treated with turbo DNase (ThermoFisher Cat. No. AM2238). To evaluate the expression patterns in the wild type and mutants; qRT-PCR primers were designed for actin, tubulin, *ELF6* and *FLC* genes (Table S7). Briefly, cDNA libraries were synthesized using Invitrogen cDNA synthesis kit (Invitrogen, Grand Island, NY) and PCR was performed using SybrGreen (Roche Cat. No. 04 707 516 001) on a LightCycler 480 (Roche, Basel Switzerland). Average C_t_ values generated from three replicates for each of the cDNA libraries were used to normalize and calculate the fold change in expression of genes, respectively.

### Reduced pod shatter phenotyping

The force required to break apart seedpods at the septum was determined using a gram force tension gauge (SSEYL ATG-100-2 Tension Gauge) attached to a two-inch alligator clip (Gardner Bender, Milwaukee, WI). Briefly, one side of a pod was clipped and then the other side was manually pulled until either the pod broke at the septum or the fin tore. Pods that broke on handling before being clipped were recorded as zero. For each line, ten pods each from five different plants were used for the measurements.

Samples for observations of separation layers were prepared by either free-hand or microtome sectioning. The free-hand sections were cut with a razor blade, briefly stained with 0.05% toluidine blue O, and then transferred to a microscope slide for observation using Nikon SMZ1500 stereo microscope. Samples used for microtome sectioning were fixed for 5 h in 2% glutaraldehyde buffered with 0.025M phosphate buffer, pH7, and post-fixed 1–2 h in 2% OsO4. Specimens were dehydrated in a graded series of acetone, infiltrated in Spurr’s resin, and polymerized in a 70° C oven. Semi-thin sections 1 µm thick were made with a Leica UCT Ultracut Microtome, stained with 0.1% toluidine blue, and viewed with a Leica DMRBE compound microscope using the 20× objective.

### Glucosinolates in leaf tissue and seeds using UV absorbance method

To determine glucosinolate content in the pennycress leaf tissues and pennycress seeds, we recorded the fresh tissue weights or seed weights of the samples. Briefly, glucosinolates were extracted in 80% methanol followed by purification and were quantified using the method described by Chopra et al. (Chopra*, et al.* 2018a). At least three biological replicates were used for estimating the glucosinolate content in each of the segregating progeny.

### HPLC analysis of wild-type and Nutty seeds

Glucosinolates in the seeds of the wild-type and mutant samples were extracted and analyzed with HPLC using the method described by Kliebenstein et al. (Kliebenstein*, et al.* 2001). Briefly, forty microliters of the glucosinolate extract was run on a 5-mm column (Lichrocart 250–4 RP18e, Hewlett-Packard, Waldbronn, Germany) on a Hewlett-Packard 1100 series HPLC. Compounds were detected at 229 nm and separated utilizing the programs described by Kliebenstein et al.(Kliebenstein*, et al.* 2001) with aqueous acetonitrile.

### Fatty acid composition analysis using Gas Chromatography

Approximately 100 mg of seeds from the wild-type and mutants of interest from NIRS scans were weighed and crushed with 1,000 µl of hexane containing a C17:0 internal standard using a mechanical homogenizer for three min at 10 m/s. The hexane supernatant containing the extracted oil was then transferred to glass vials for methylation and FAMES were separated and detected using the methods described in Chopra et al (Chopra*, et al.* 2018a). At least two replicates were used for estimating the fatty acid composition in each of the lines.

## General

We acknowledge the hard work of many undergraduates and others who contributed to this study including: Karl Nord, Carl Branch, Liam Sullivan, Luke Aldrich, Greta Rockstad and many others. We thank Martha Cook and Tara Nazarenus for help with tissue embedding / sectioning / microscopy and seed oil analyses, respectively.

## Funding

This material is based upon work that is supported by the National Institute of Food and Agriculture, U.S. Department of Agriculture, under award numbers 2014-67009-22305 and 2018-67009-27374 to M.D.M, J.C.S and W.B.P. Funding for research conducted in the E.B.C lab was provided by U.S. Department of Energy, Office of Science, OBER (DOE-BER SC0012459) and NSF Plant Genome Program (13-39385). Additional funds were provided by the Minnesota Department of Agriculture and the University of Minnesota Forever Green Initiative to J.A.A. and B.I., and the Walton Family Foundation, PepsiCo and General Mills to D.L.W.

## Author Contributions

M.D.M. and J.C.S. conceived the study, designed experiments, supervised and organized coworkers, created mutagenized populations, isolated mutants, helped characterize the mutants, identified candidate genes, and co-wrote the manuscript. R.C. along with E.B.C, C.G. and B.I. characterized the fatty acid profiles, along with K.M.D. performed WGS analyses, and also independently performed extensive data analyses and helped write the first draft of the manuscript. J.L. D.J.K, and T.U. were responsible for the wet lab glucosinolate analyses. E.D., N.F, R.E, M.E., M.M., A.B. and K.A. isolated mutants and helped characterize candidate genes. M.O. helped evaluate *FLC* expression and seed yield of the *elf6* mutant. J.A.A. and K.A. help with the initial NIR set-up. K.M.D, K.F. and R.C. helped characterize the *elf6* mutant. A.B. generated and imaged wild-type and *ind-3* seedpod sections. D.L.W. isolated wild-type line MN106 used in the study and aided in planning.

## Competing Interests

The authors declare potential competing financial interests as intellectual property applications have been submitted on portions of the reported research.

## Data and materials availability

All sequence information described in this study is contained within the Supplementary Materials. All plant materials described in this report are available upon completion of Material Transfer Agreements.

## Supplementary Materials

### List

**Figure S1.**
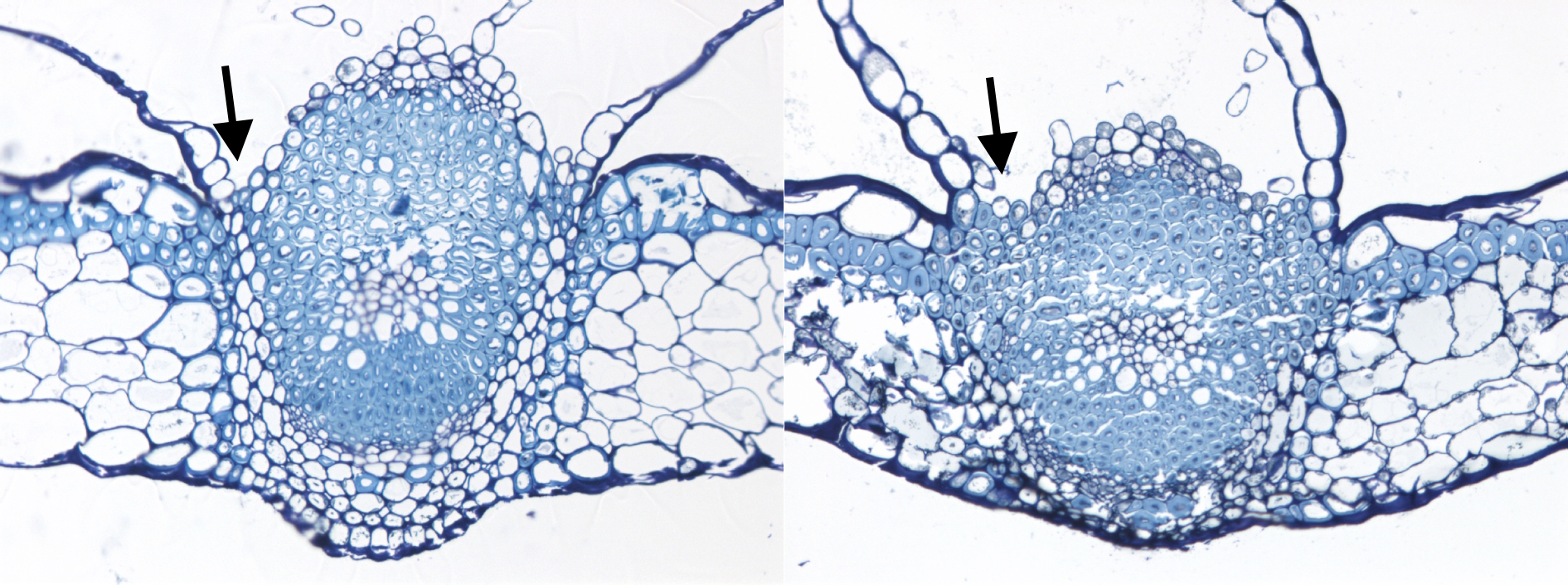
Comparisons of the separation layer of wild-type Spring 32 (left) and *ind-3* (right) seedpod sections stained with toluidine blue. Note the lack of a clear separation layer in *ind-3* (arrows).

**Figure S2:**
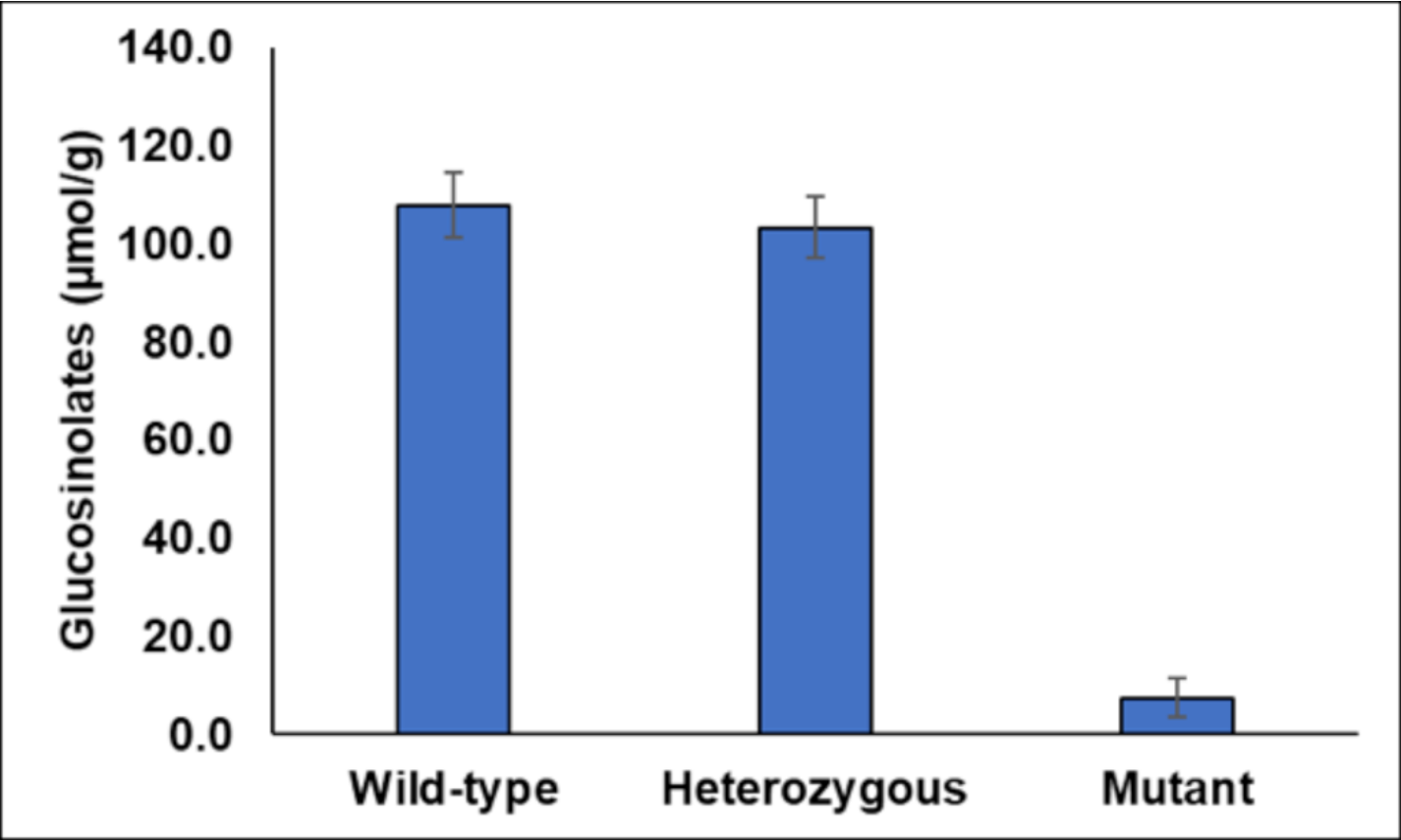
Average glucosinolate content of the progenies from the segregating population of *Ta-aop2-1* plants. Allele-specific markers were used to assess the *aop2-1* genotype of members in the population and glucosinolate content was analyzed using NIRS and confirmed with wet-lab assay.

**Figure S3.**
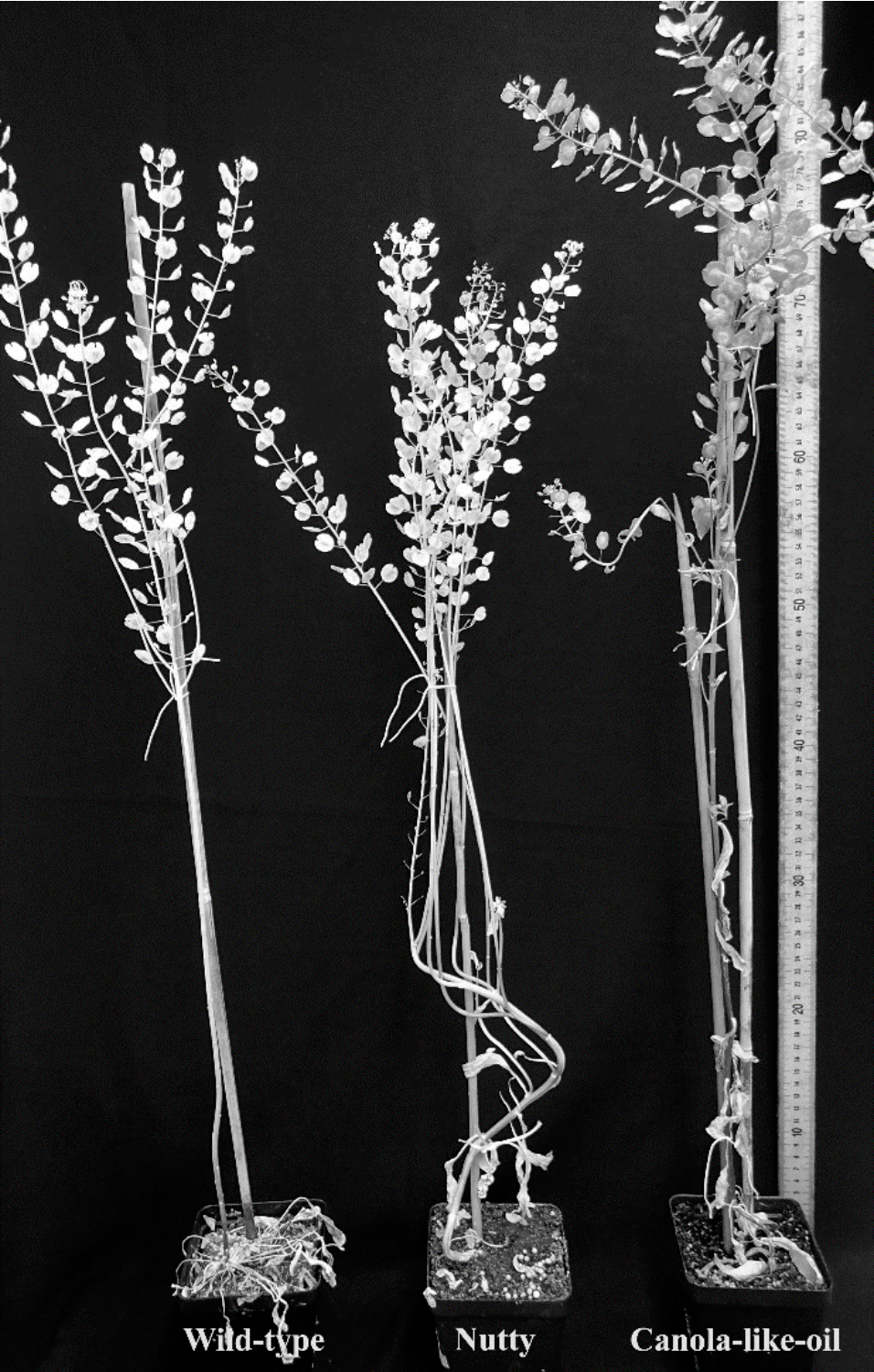
Comparisons of a wild-type pennycress plant with the “Nutty” *aop2-1* and “canola-like-oil” *fae1-1 rod1-1* mutants grown in controlled chambers. No developmental defects were observed to be associated with the respective traits.

**Figure S4.**
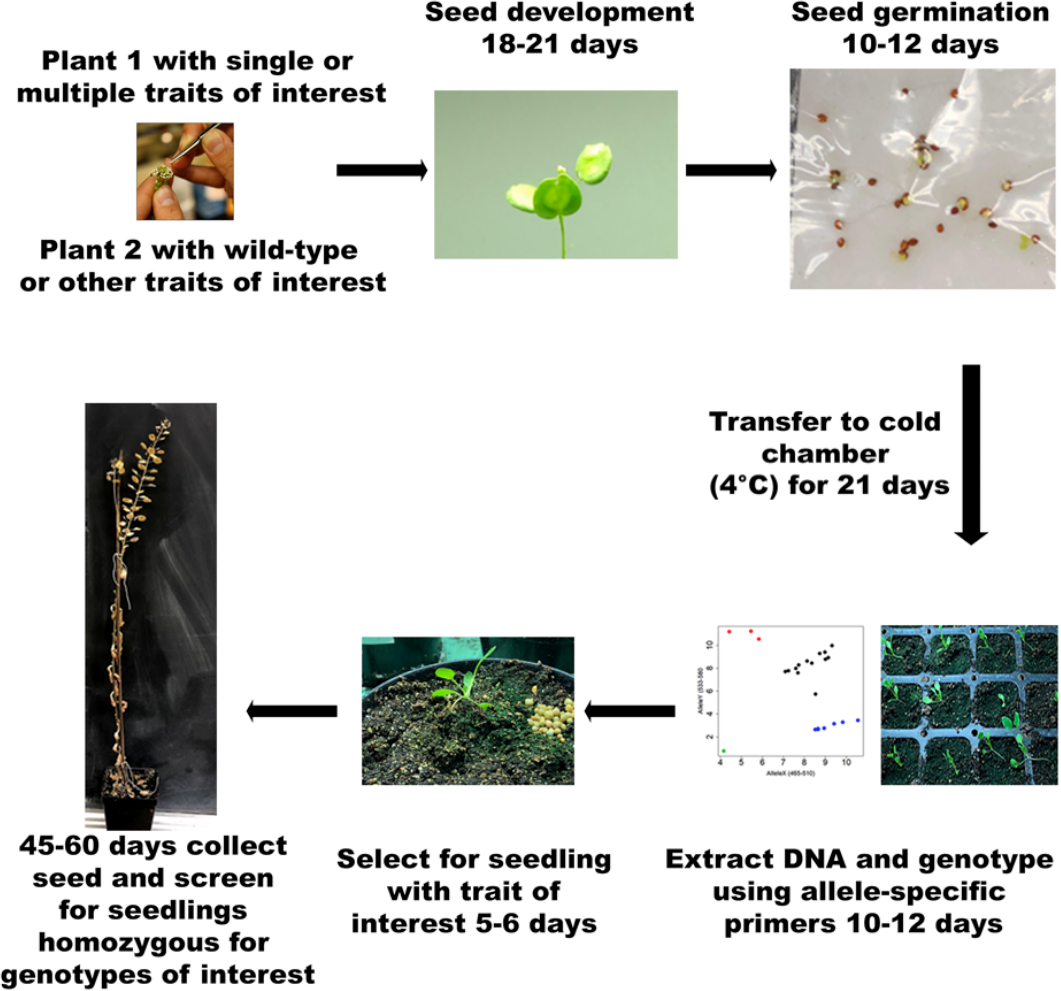
A schematic representation of pennycress life-cycle from seeds to a mature plant. Seeds can be developed from the crossing event in 12-15 days. Seeds can be propagated from seeds to mature in 75-80 days. This scheme of propagating pennycress from would allow for minimum of three generations per year.

**Figure S5.**
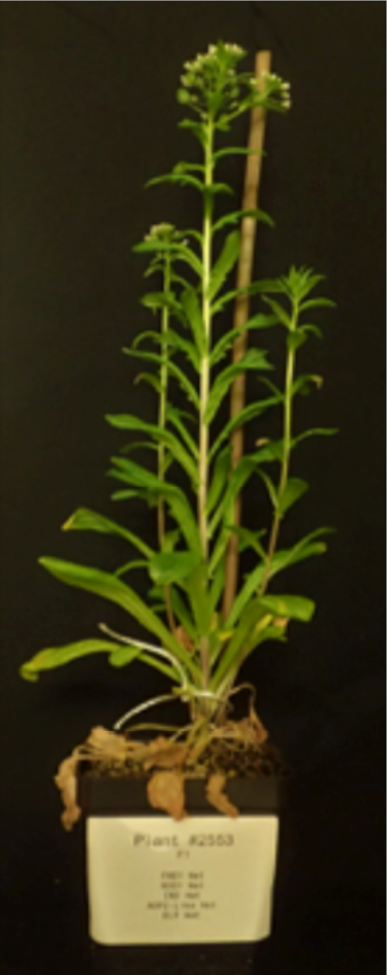
F_1_ plant carrying all of the described mutations in this report and was selected using the allele-specific primers and strategy described in Fig S4.

**Table S1:** Plant height and seed yield comparisons between wild-type and early flowering mutant (elf6-1) grown in 1 × 1 m plots in the field. No statistically significant differences were observed among these lines (± sd).

| Line | Plant height (cm) | Seed yield (g) |
| --- | --- | --- |
| Wild-type | 65 ($\pm 0.25$) | 46.56 ($\pm 11.48$) |
| elf6-1 | 67.58 ($\pm 0.76$) | 57.61 ($\pm 14.15$) |

**Table S2.** Protein domain analysis to highlight the conserved region in the Indehiscent gene (IND) using several databases.

| Gene | Database | Family number | Start Pos. | End Pos. | Gene Family |
| --- | --- | --- | --- | --- | --- |
| Ta1.0_25465 | SMART | SM00353 | 98 | 147 | Myc-type, basic helix-loop-helix (bHLH) domain |
| Ta1.0_25465 | Gene3D | G3DSA:4.10.280.10 | 84 | 154 | Helix-loop-helix DNA-binding domain superfamily |
| Ta1.0_25465 | ProSiteProfiles | PS50888 | 92 | 141 | Myc-type, basic helix-loop-helix (bHLH) domain |
| Ta1.0_25465 | SUPERFAMILY | SSF47459 | 97 | 151 | Helix-loop-helix DNA-binding domain superfamily |
| Ta1.0_25465 | CDD | cd00083 | 96 | 146 | Myc-type, basic helix-loop-helix (bHLH) domain |
| Ta1.0_25465 | Pfam | PF00010 | 100 | 141 | Myc-type, basic helix-loop-helix (bHLH) domain |

**Table S3.** Summary of the mutations in the domestication related genes characterized in this study.

| Scaffold Number | Gene | # of Alleles | Mutation |
| --- | --- | --- | --- |
| Ta_scaffold_10 | *ELF6*<br>Early Flowering6 | 1 | **Mutant:** A7 25<br>**Genomic pos:** 107096<br>**cDNA pos:** C952T<br>**AA pos:** His318Tyr |
| Ta_scaffold_1003 | *IND*<br>Indehiscent | 3 | **Mutants:** E5 552; E5 550; Spring 32 NS<br>**Genomic pos:** 6741, 6754, 6795<br>**cDNA pos:** C247T; G260A; G301A<br>**AA pos:** Pro83Ser; Arg87His; Ala101Thr |
| Ta_scaffold_74 | *AOP2*<br>Alkenyl- and Hydroxyalkyl-Producing2 | 1 | **Mutant:** E3196<br>**Genomic pos:** 395125<br>**cDNA pos:** G289A<br>**AA pos:** Gly97Arg |
| Ta_scaffold_45 | *FAE1*<br>Fatty Acid Elongation1 | 2 | **Mutants:** V296; K1822<br>**Genomic pos:** 254962;<br>**cDNA pos:** C1018T; G1349A<br>**AA pos:** Gln340Stop; Trp450Stop |
| Ta_scaffold_199 | *ROD1*<br>Reduced Oleate Desaturation1 | 2 | **Mutants:** d0422; E5 370 P6<br>**Genomic pos:** 186474; 186409<br>**cDNA pos:** G678A; G743A<br>**AA pos:** Met226Ile; Arg248Lys |

**Table S4.**
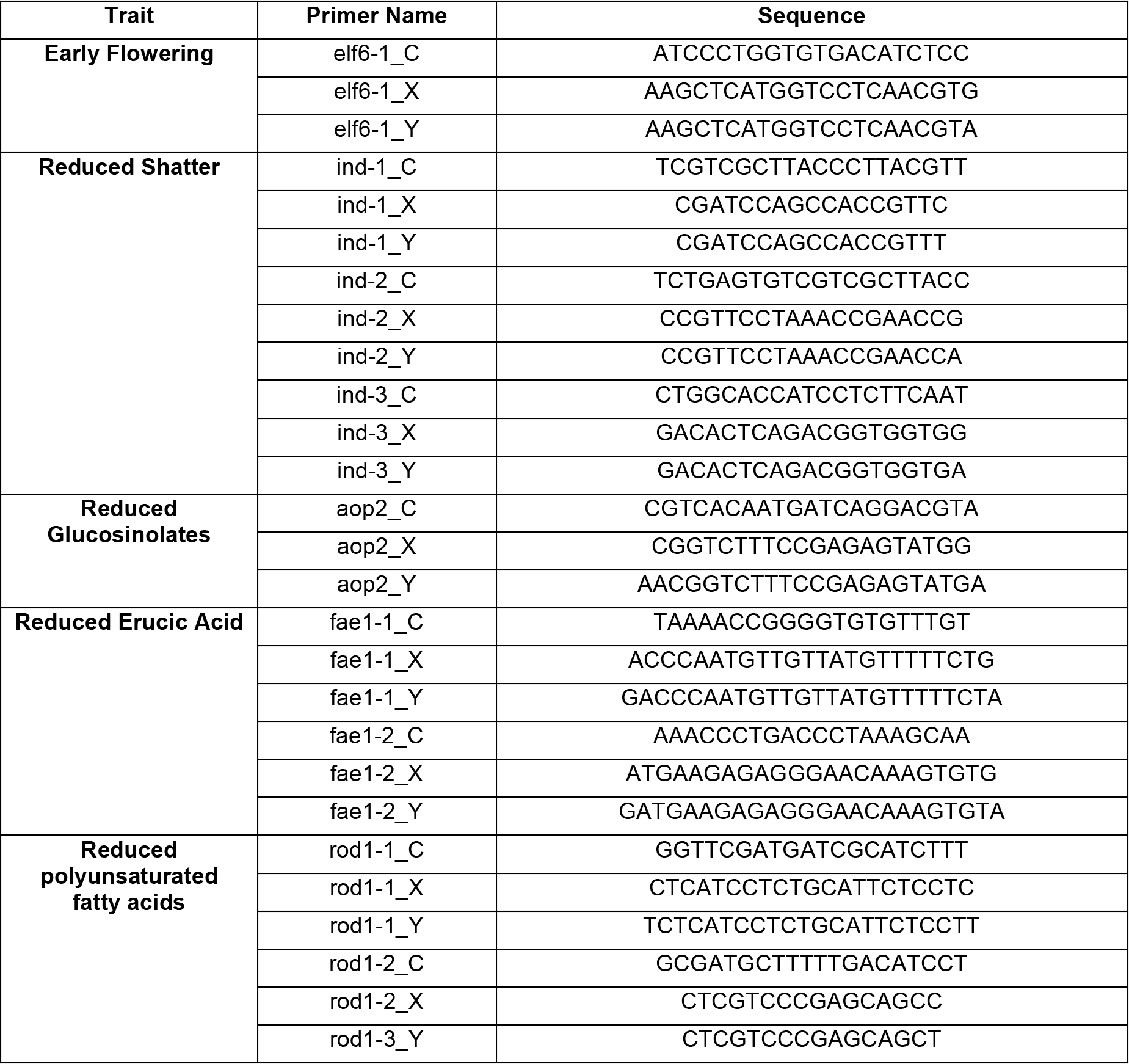
Allele-specific primers designed for single nucleotide mutations in the domestication candidate genes.

**Table S5:**
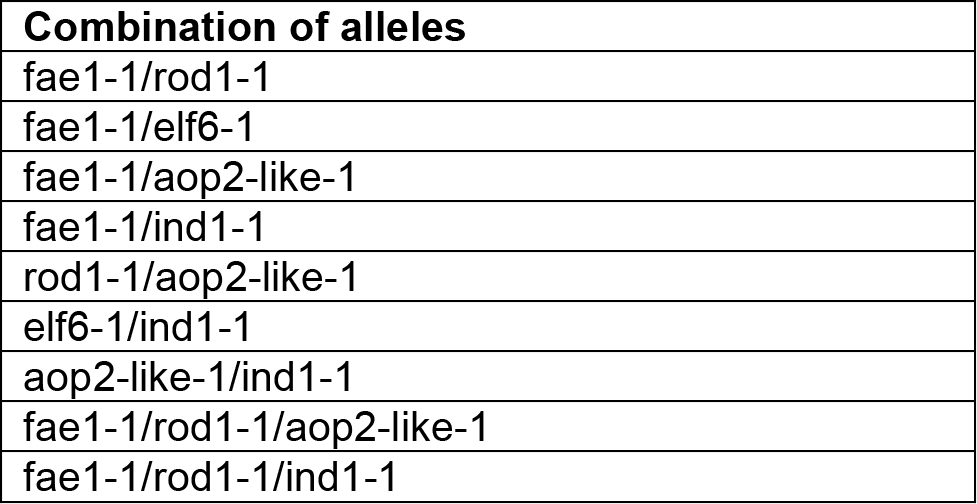
Summary of double and triple mutants generated using the approach described in Fig S4.

**Table S6.**
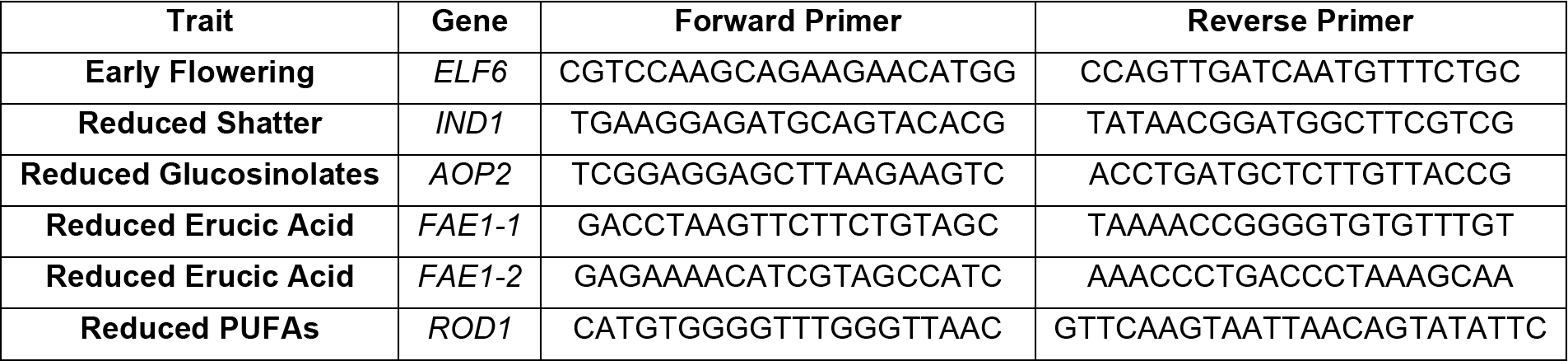
Primers used for Sanger sequencing.

**Table S7.**
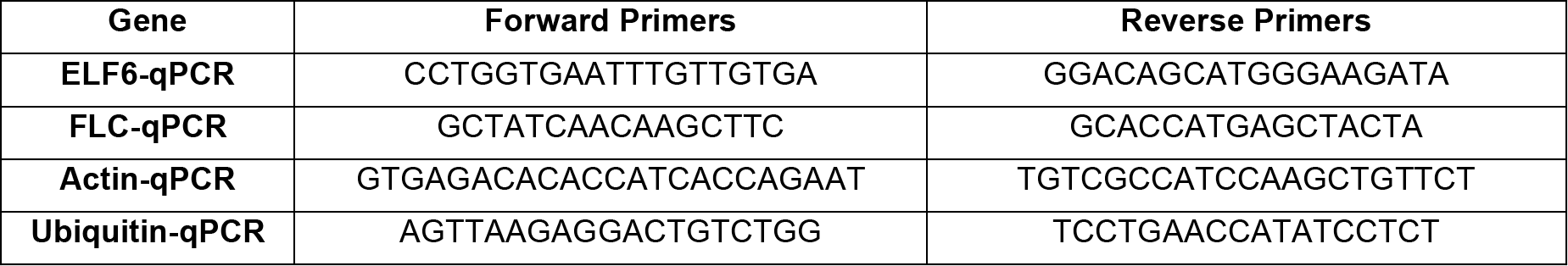
Primers used for expression analysis of the *Ta-elf6-1* early flowering mutant.

**Data S1.**
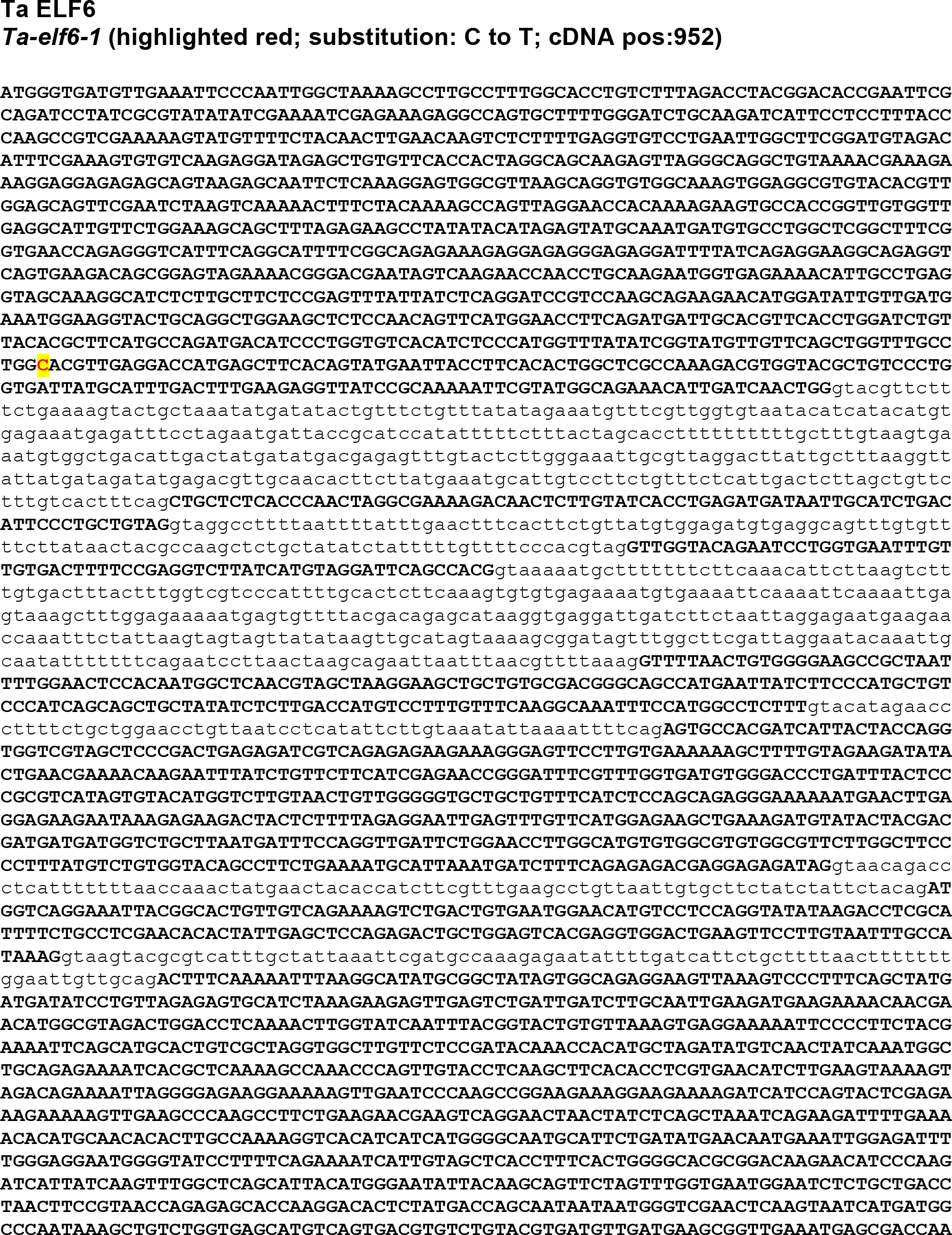
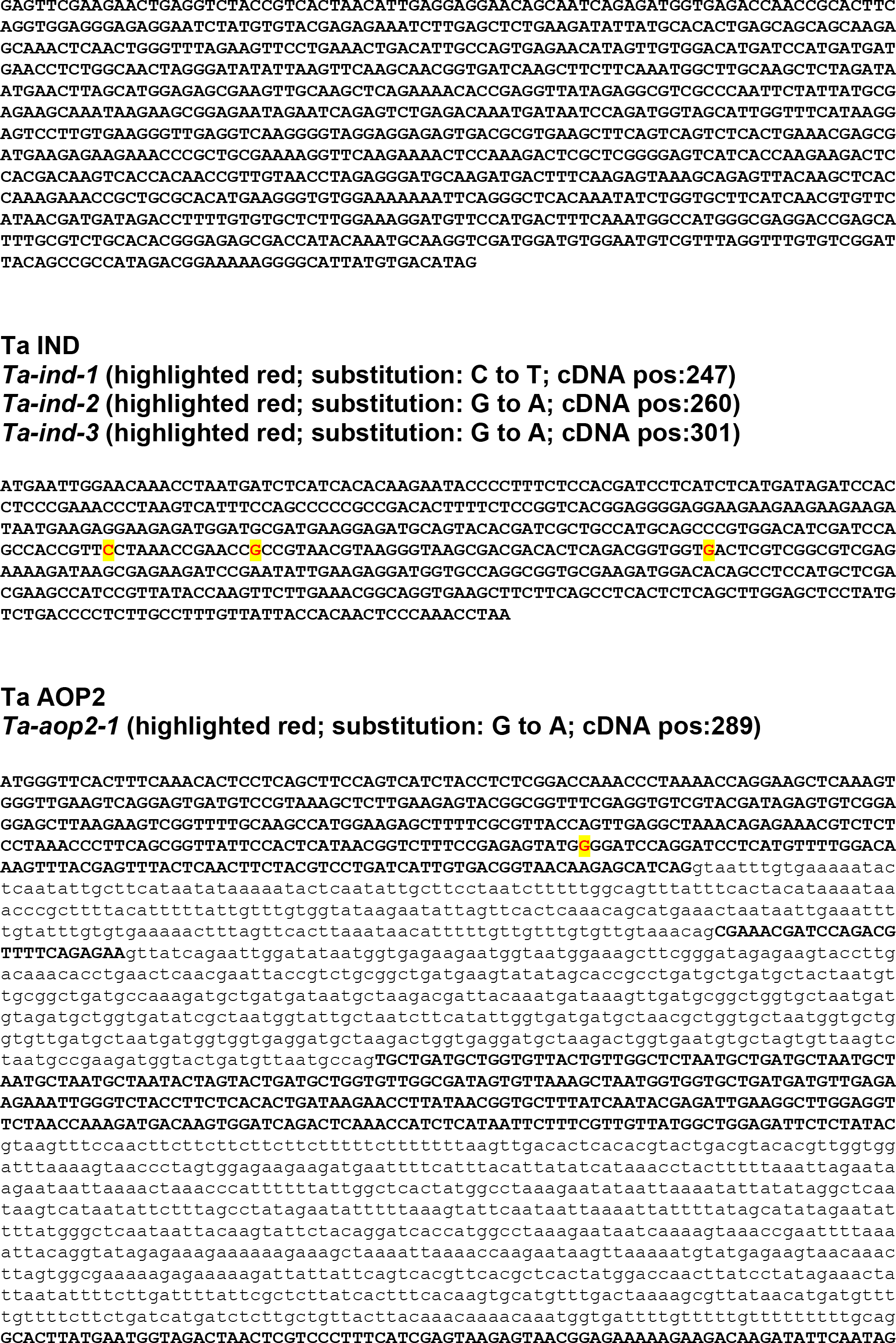
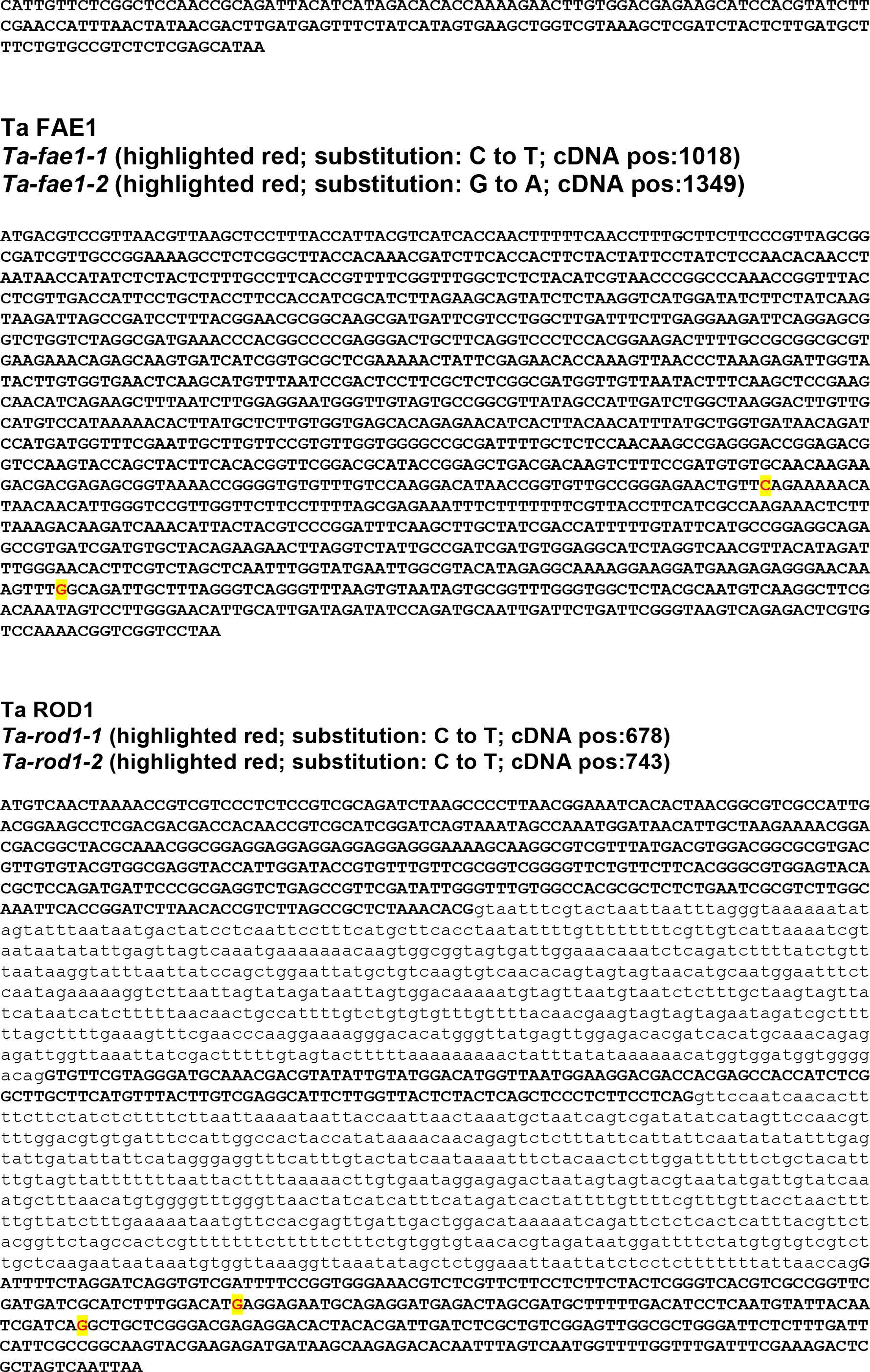
Genomic DNA Sequences of the candidate genes for pennycress domestication obtained from Thlaspi (pennycress) version 1.0 gene annotations and information on the mutation sites. (Exons – bold; Introns – small letters).

**Data S2.**
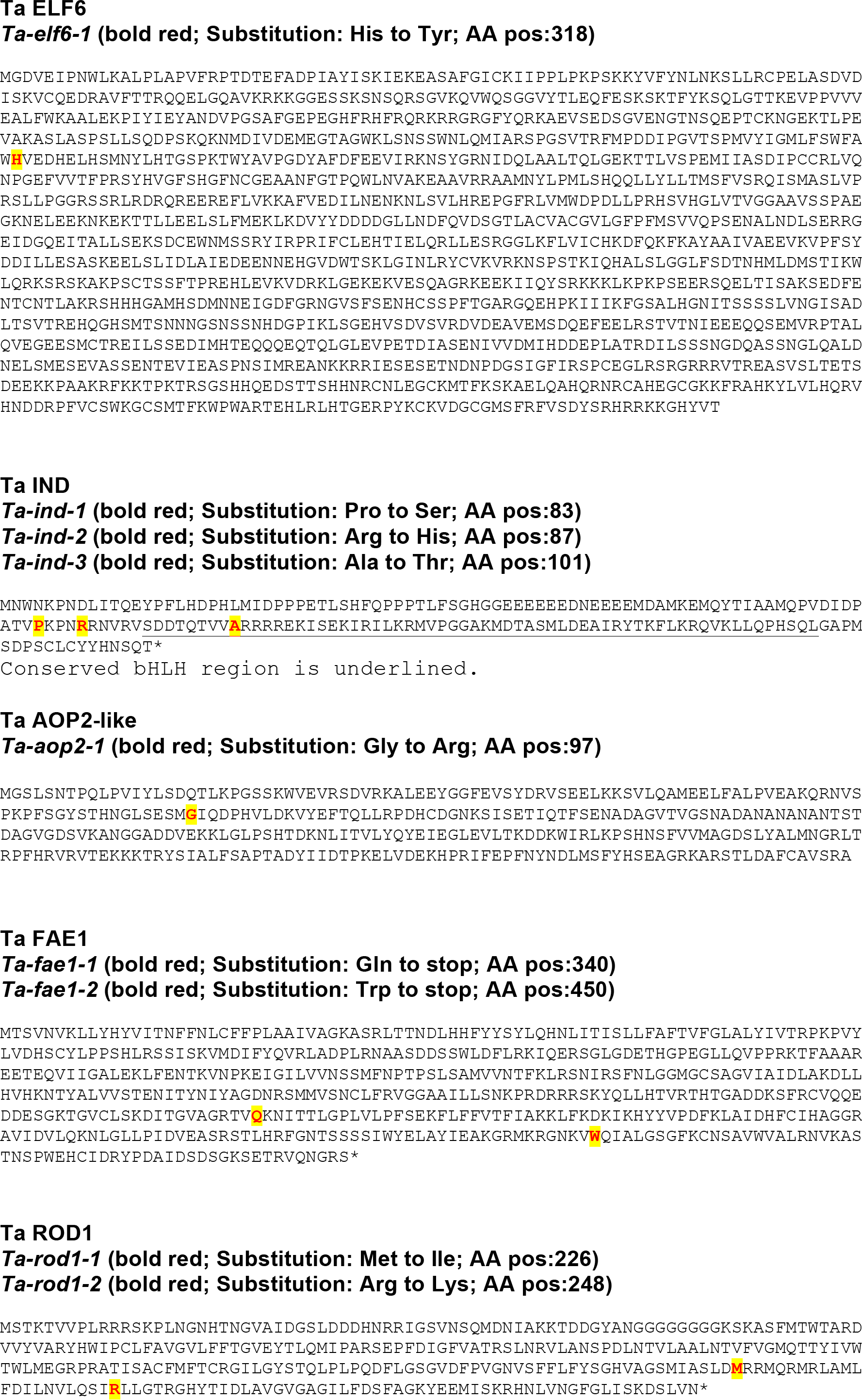
Predicted protein sequences of the candidate genes for pennycress domestication obtained from Thlaspi version 1.0 gene annotations and mutation and information on the mutation sites.

**Data S3.**
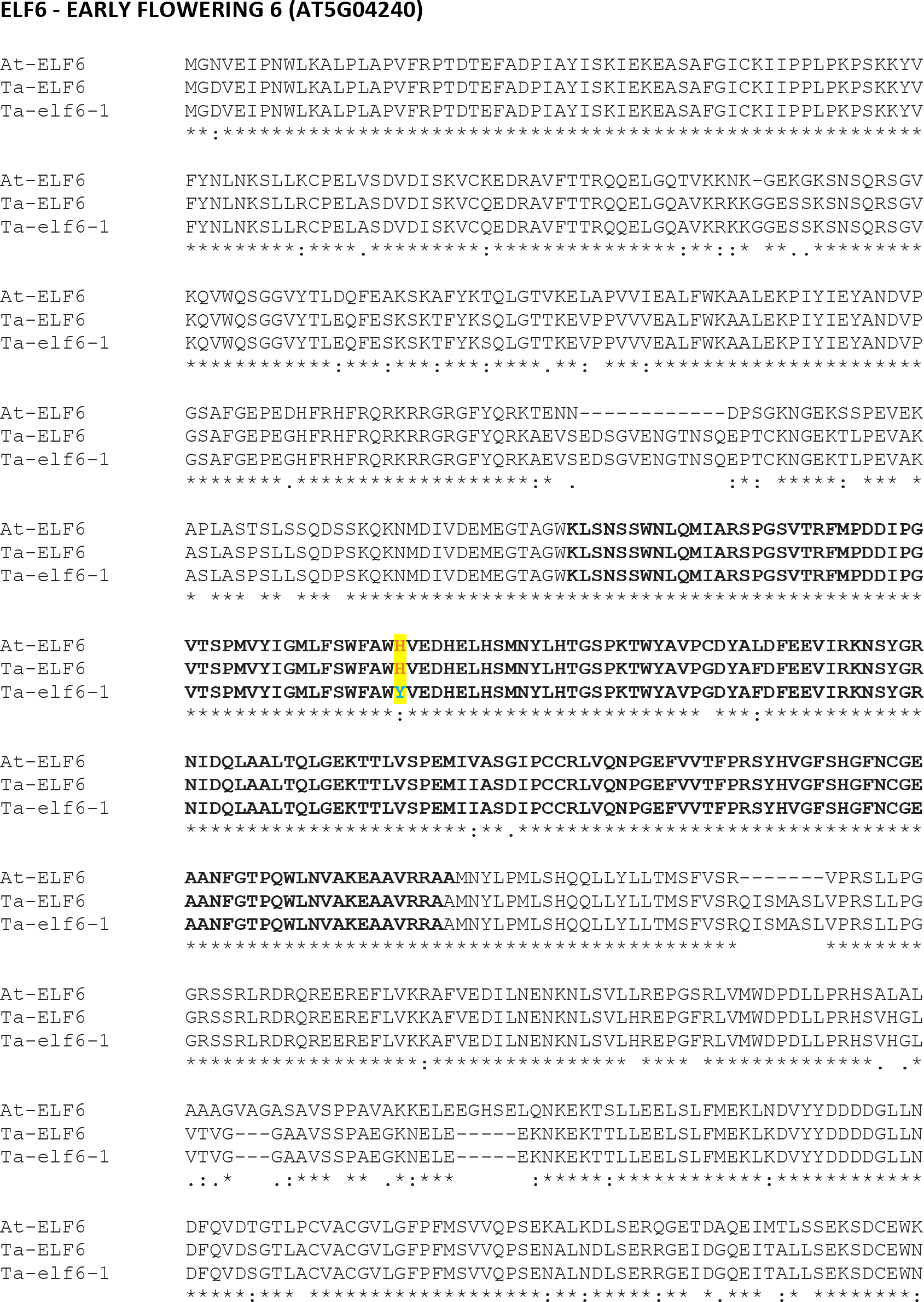
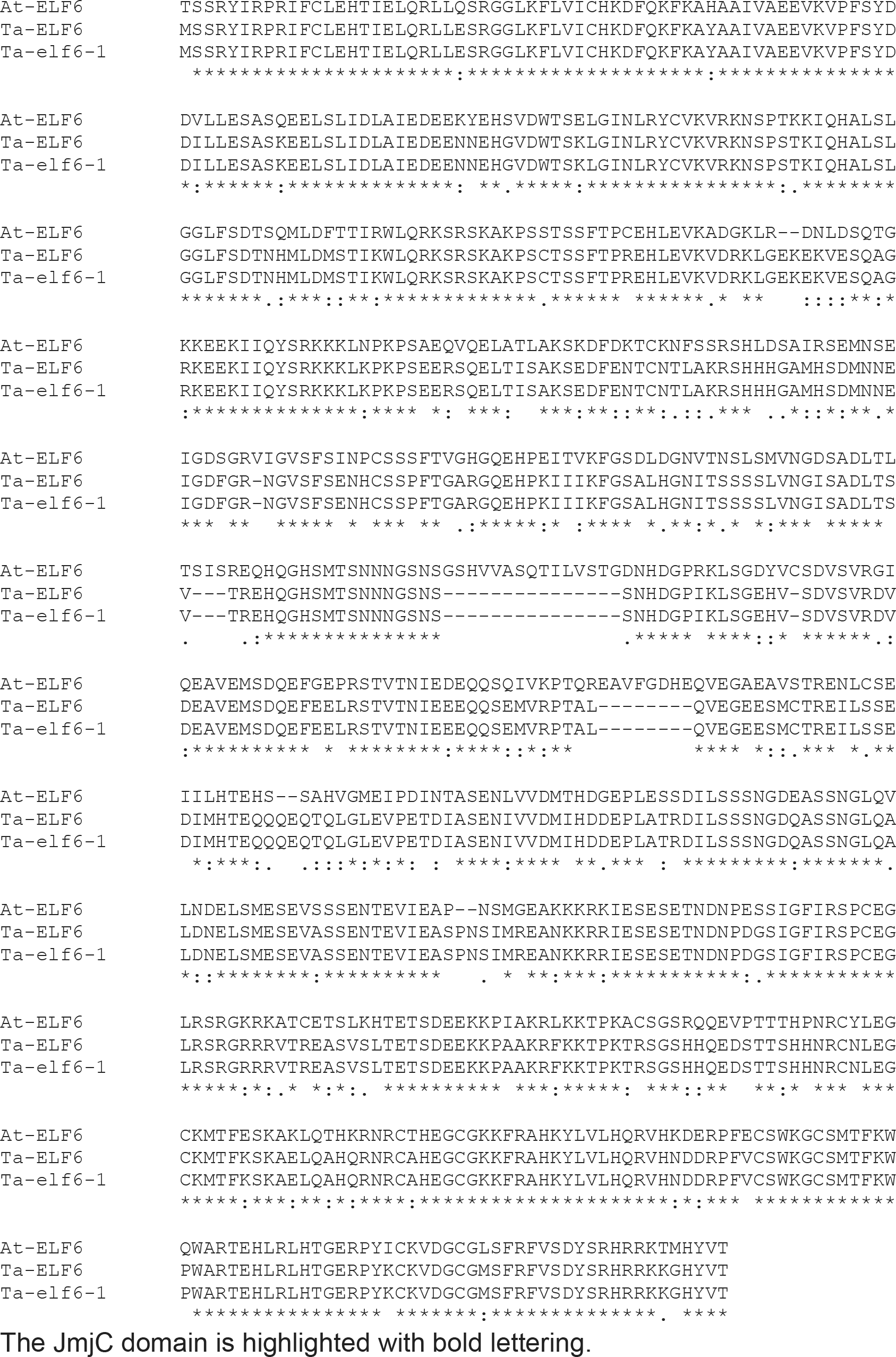
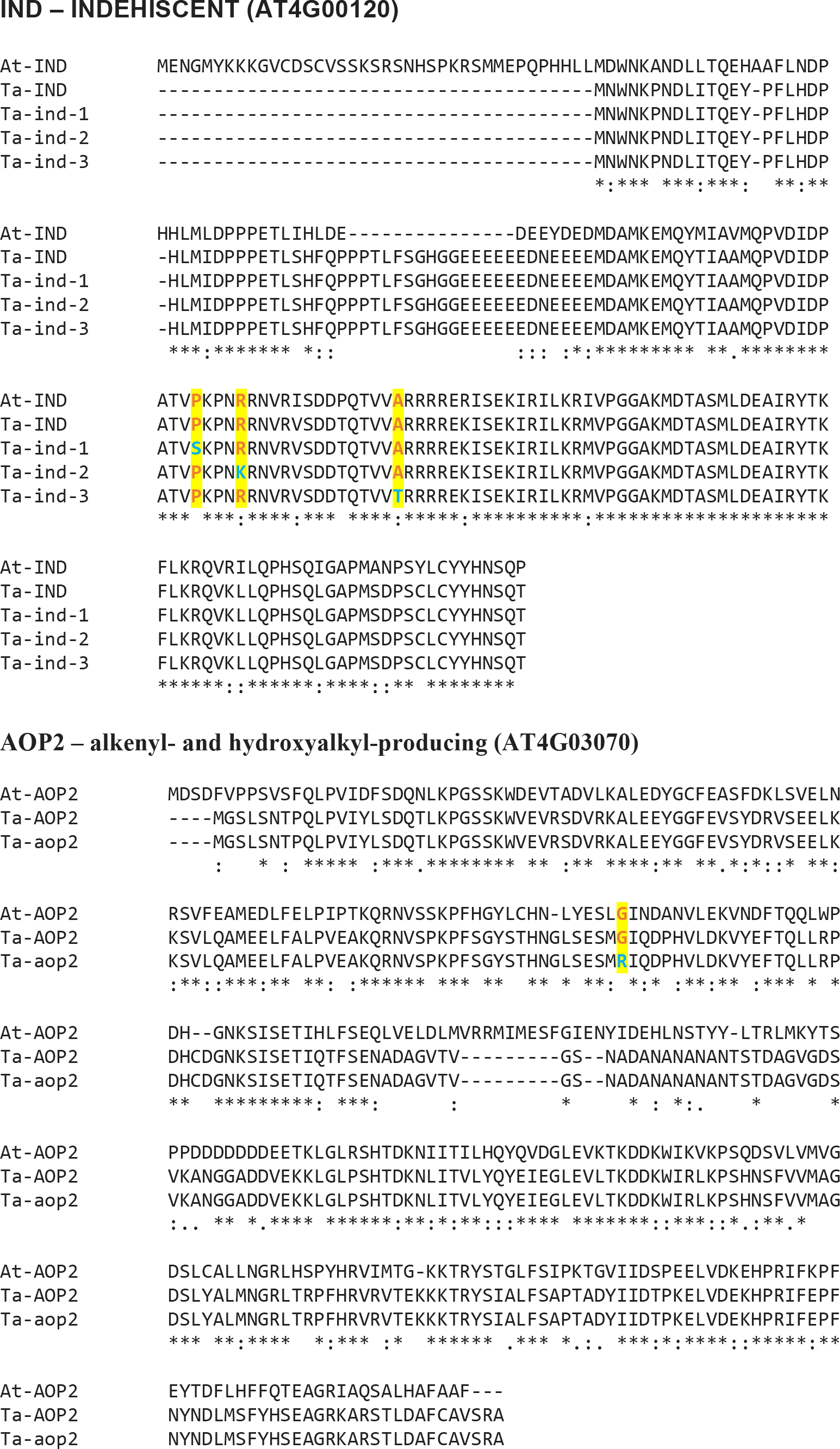
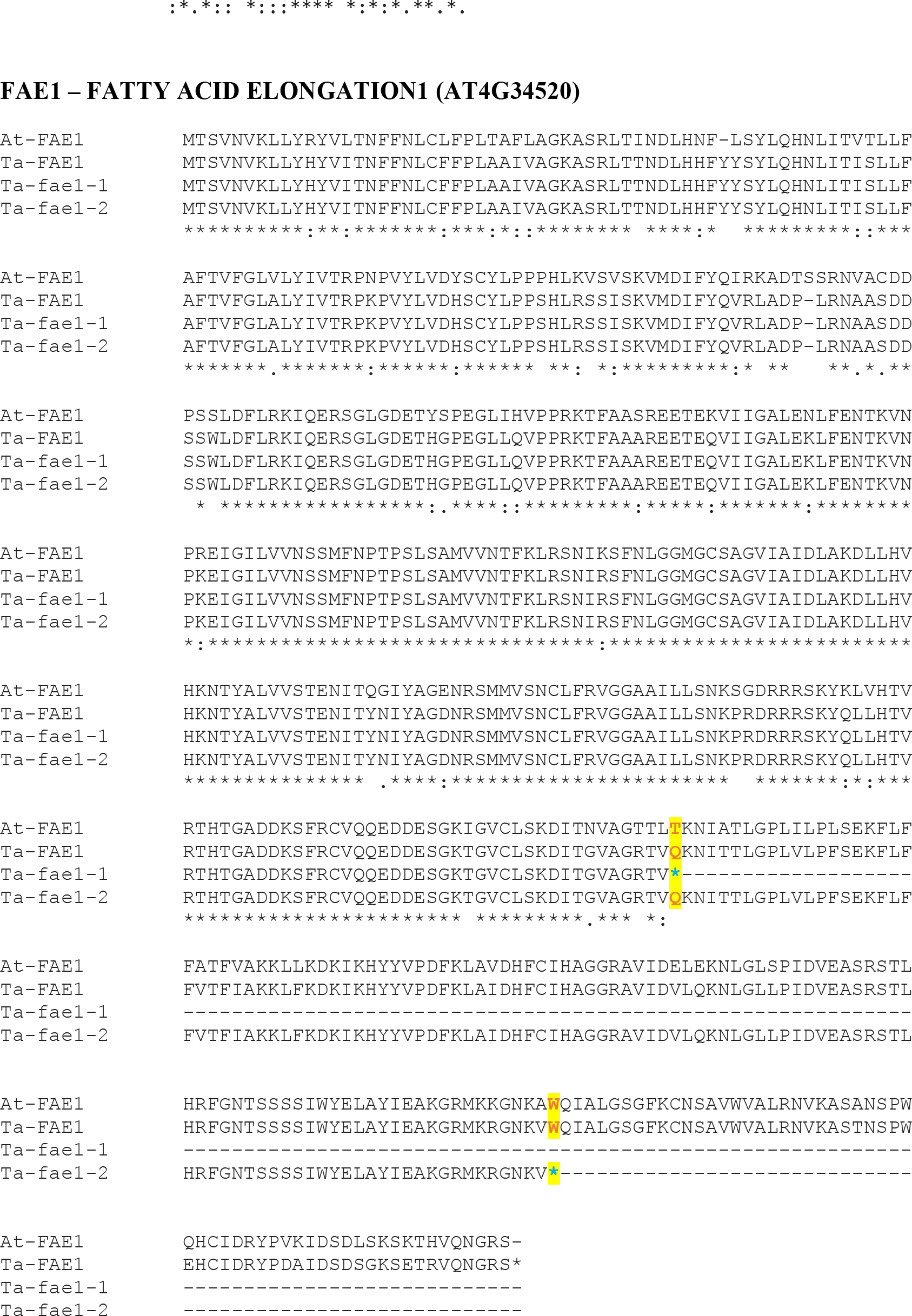
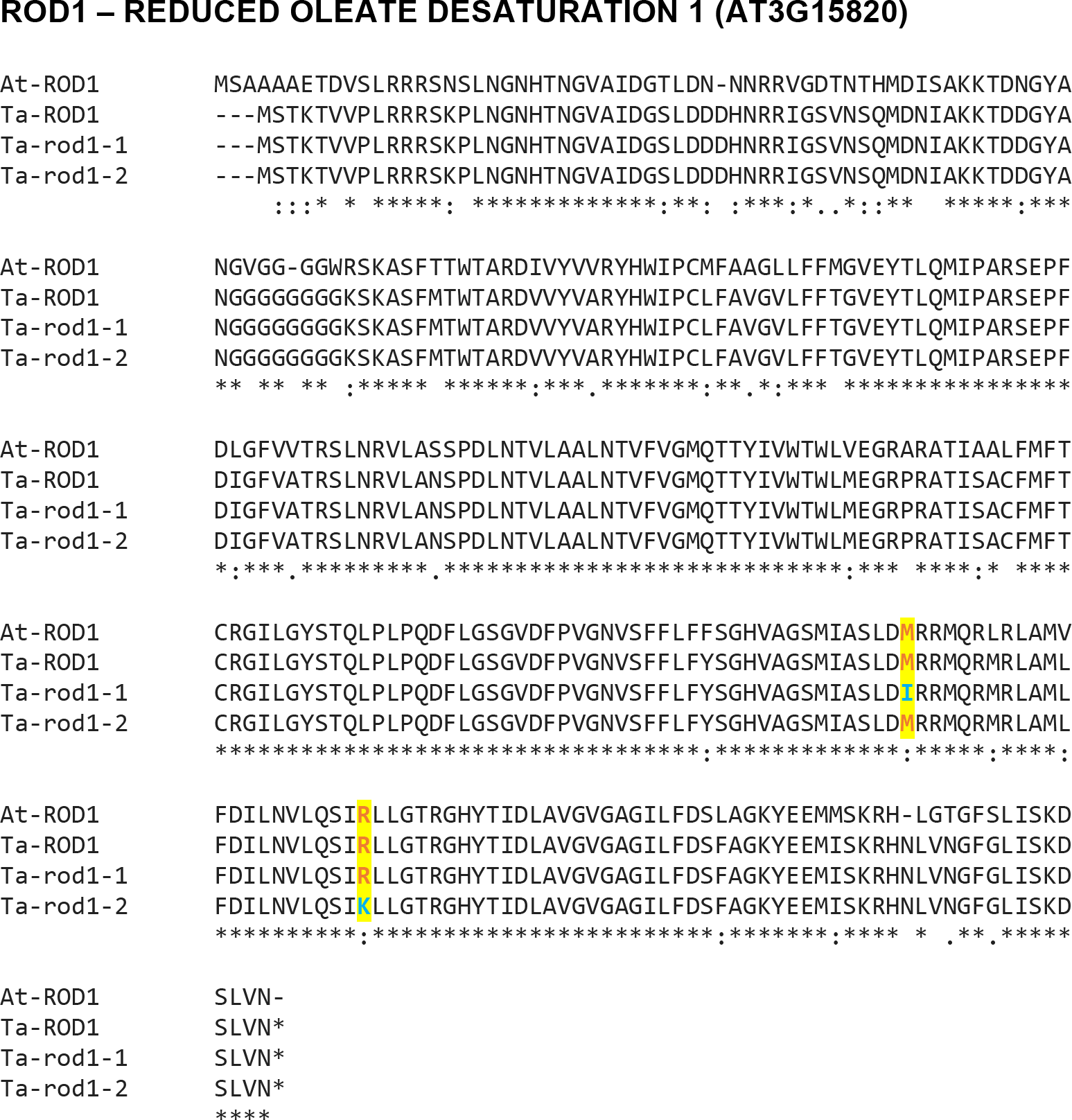
Protein sequence alignments of *Thlaspi arvense* (pennycress) wild-type sequences with the corresponding mutant and orthologous Arabidopsis sequences. At the locations of the mutations, wild-type amino acids are highlighted in red and mutant amino acids in blue.

